# Real-time reorienting during episodic retrieval rescues memories from suboptimal arousal-attentional states

**DOI:** 10.64898/2026.10.28.762693

**Authors:** Shawn T. Schwartz, Haopei Yang, Mingjian He, Khanh K. Nguyen, Megan T. deBettencourt, Tammy T. Tran, Kevin P. Madore, Anthony D. Wagner

**Affiliations:** Department of Psychology, Stanford University; Wu Tsai Neurosciences Institute, Stanford University; Stanford University Wu Tsai Human Performance Alliance; Stanford Center for Mind, Brain, Computation and Technology; Department of Anesthesiology, Perioperative, & Pain Medicine, Stanford Medicine; Department of Psychological & Brain Sciences, Washington University in St. Louis; School of Psychology, Georgia Institute of Technology; Johnson & Johnson, USA

**Author notes:** **Correspondence** should be addressed to S.T.S.

## Abstract

Successful knowledge expression varies with one’s arousal-attentional (A-A) state at the moment just prior to attempting to remember. Yet, whether such states causally affect retrieval remains unclear. Here we introduce PRISM (Pupil-based Real-time Intervention System for Memory), a closed-loop paradigm that continuously monitors pupil size to track A-A states in real time and, during suboptimal states (SOS), delivers a visual reorienting cue in the moment before a retrieval attempt. In 67 participants performing a goal-directed source memory and item recognition task, we show that pupil-defined SOS just prior to retrieval impair both item recognition and source recollection relative to optimal state (OS) trials. Critically, delivering a brief visual reorienting trigger cue on SOS trials in real time rescues item memory, but not source memory, raising item memory to a level not credibly different from real-time OS trials and providing causal evidence that phasic attentional reorienting during A-A-defined suboptimal states facilitates memory access at retrieval. Offline spline-detrending analysis confirms that the A-A state-memory relationship is robust to slow physiological drift. It also shows that the trigger benefit was credible for marginally suboptimal (drift-driven) SOS trials and smaller and less certain for deeply suboptimal (drift-resistant) SOS trials, consistent with a zone-of-recoverability account, although the difference between the two SOS subtypes was not credible. Furthermore, among deeply suboptimal trials the A-A-boosting trigger cue improved memory for low-A-A SOS (i.e., pupillary constriction) but not high-A-A SOS (i.e., pupillary dilation) trials, consistent with the classic inverted-U relationship between arousal and performance, although the difference between the two directions was not credible. Together, these findings causally link phasic arousal-attentional reorienting at retrieval to memory rescue and suggest that the depth and direction of suboptimality shape whether such rescue succeeds, connecting the readiness-to-remember framework to the classic arousal-performance inverted-U relationship. Moreover, these findings provide a further foundation for adaptive closed-loop technologies that would tailor intervention to the depth and direction of momentary arousal-attentional states, facilitating memory expression in real time.

## Introduction

Moment-to-moment fluctuations in preparatory attention and arousal explain, in part, why certain moments are more conducive to successful memory than others [1–14]. Evidence from neuroimaging, electrophysiology, and pupillometry indicates that such fluctuations reflect systematic shifts in the brain’s arousal and attentional states, henceforth, ‘A-A states’ [15–23]. Correlations between read-outs of A-A states, cognitive control, and memory motivate readiness-to-learn and readiness-to-remember frameworks [5, 6, 14, 24–28], which build on a rich literature linking preparatory states to cognitive performance [27, 29–42]. Notably, the apparent effects of A-A states at the time of retrieval highlight fundamental challenges for education, clinical populations with attention dysregulation, and everyday cognitive performance: well-encoded memories are not sufficient for performance if attention and arousal fluctuations in the moment just prior to an attempt to remember decrease the probability of successful memory retrieval.

The adaptive gain theory of locus coeruleus (LC) function suggests that the relationship between tonic arousal/attention and cognitive performance follows an inverted-U shaped curve [43–45]. Pupil size, a non-invasive and continuous proxy index of LC-norepinephrine (NE) activity, tracks fluctuations in arousal and attention with relatively high temporal resolution [46–50]. Both tails of the pupil size distribution have been linked with distinct modes of attentional failure. Relatively small pre-stimulus pupil size may reflect a hypoactive LC mode characteristic of internally-oriented inattention and mind wandering, whereas comparatively larger pre-stimulus pupil size might index a hyperactive LC mode associated with external distraction and exploratory disengagement [47, 51]. Greater variability in pre-stimulus pupil size has likewise been related to poorer performance both within and across participants [47, 48, 52], consistent with the view that frequent departures away from an intermediate arousal optimum can be detrimental to sustained cognitive engagement and performance. During cognitively demanding task sessions, pre-stimulus pupil fluctuations give a useful proxy for variation of arousal/attention. Moreover, relative deviations in pre-stimulus pupil size are associated with meaningful variation in memory performance [5, 6, 17, 28, 53–58]. In humans, a wealth of evidence documents how A-A processes and states during the act of retrieval itself can impact memory outcomes [59–66], with complementary evidence from animal models and pharmacological studies suggesting that the LC-NE system plays an important role during memory retrieval such as norepinephrine-modulated hippocampal pattern completion [67, 68].

More recently, the field has focused on measuring and understanding the preparatory mechanisms that immediately precede acts of retrieval and correlate with memory outcomes (i.e., readiness-to-remember). These include read-outs of arousal, attention, and/or goal state processes just prior to and during expression of episodic memories [1–6, 14, 69]. A primary aim of the present study is to complement traditional correlational approaches to characterizing the relationship between preparatory A-A states and retrieval performance, using a closed-loop approach to garner causal evidence for the role of A-A states in episodic retrieval. To do so, we developed the PRISM (Pupil-based Real-time Intervention System for Memory) paradigm, a closed-loop pupillometry system that enables real-time monitoring of and intervention on A-A states at retrieval. PRISM continuously monitors raw pupil size — an index of tonic LC-NE A-A state — in the moments just prior to each memory retrieval trial, detects A-A-defined suboptimal states (SOS) in real time via fluctuations in tonic pupil within a block, and, on a subset of trials, triggers delivery of a visual reorienting cue designed to phasically perturb the A-A state before the retrieval probe appears (Fig 1c). This design creates a within-subject, within-session comparison between SOS trials that did not receive intervention and optimal state (OS) trials and between SOS trials that did versus did not receive intervention, enabling causal inference about the role of pre-retrieval A-A state in memory retrieval performance. In addition, by leveraging detected suboptimal states with different signs in their pupil deviation, we can examine predictions from the inverted-U framework, which may differ in their amenability to recovery by a visual reorienting cue.

**Fig 1.**
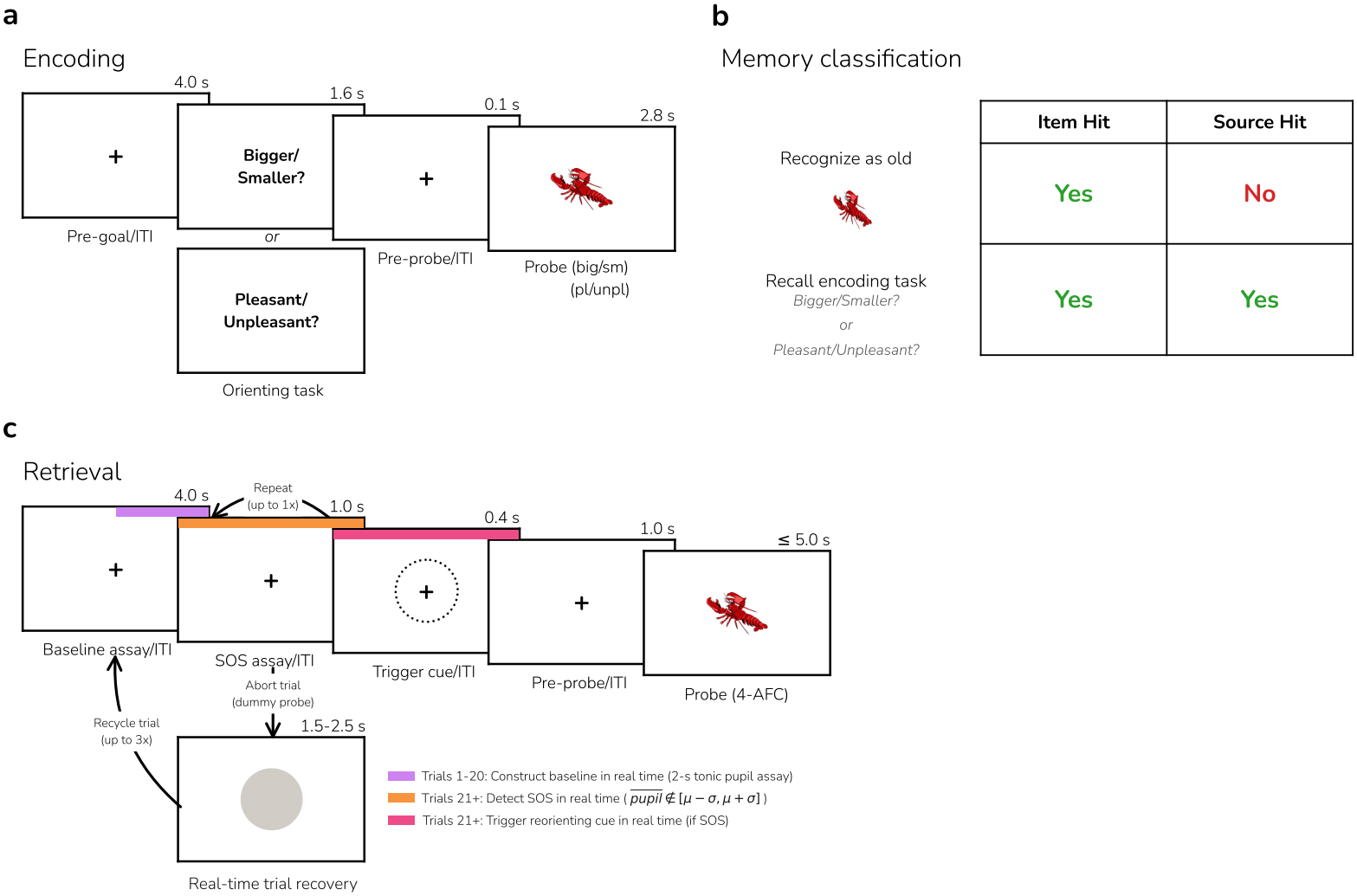
PRISM’s real-time experimental design. (**a**) Participants viewed 168 objects once across 3 intentional encoding blocks of 56 trials each. In each encoding block, 28 objects were classified on a conceptual goal dimension (pleasant or unpleasant) and 28 on a perceptual dimension (bigger or smaller), yielding 84 conceptual object trials and 84 perceptual object trials across blocks. Each object appeared either bigger (450 *×* 450 px) or smaller (150 *×* 150 px) in size on the screen, independent of goal cue, counterbalanced across participants through random assignment. (**b**) Definitions of item hit and source hit memory outcome categories for studied objects. (**c**) After each encoding block, participants performed a retrieval block for the just-encoded objects, wherein they viewed 84 objects—56 studied and 28 novel foils. On each trial, participants made a four-alternative forced-choice (4-AFC) memory judgment on a medium-sized object (300 *×* 300 px), indicating whether they: (1) remember performing the size decision (i.e., bigger or smaller before) with the object, (2) remember performing the pleasantness decision (i.e., pleasant or unpleasant before) with the object, (3) remember encountering the object during encoding but cannot remember which decision was performed, or (4) do not remember encountering the object and thus perceive it as new. Critically, beginning on trial 21 of each test block, real-time pupillometry was conducted across either one or two 1-s sampling windows to assess momentary pupil dilation. If the mean pupil assay during the sampling window exceeded a threshold of *±*1 s.d. from the fixed baseline mean (computed across the first 20 trials of the block), the A-A state was deemed suboptimal, and a reorienting cue was delivered on one randomly chosen trial of each successive pair of SOS trials.

We hypothesized that real-time triggering on SOS trials prior to retrieval will yield better memory performance than untriggered SOS trials, providing causal evidence that phasic perturbation during A-A-defined suboptimal states at retrieval rescues memory access as consistent with the readiness-to-remember framework [5, 6, 14]. Moreover, because pupil-defined SOS trials can reflect either abnormally low arousal (pupil constriction) or abnormally high arousal (pupil dilation), and because the intervention is a brief alerting stimulus that we predicted would boost phasic arousal, we further hypothesized that the theoretical inverted-U relationship between arousal and performance [45] would result in an asymmetric influence of the intervention. Specifically, we hypothesized that the intervention would be effective for low-arousal SOS trials, which can be nudged toward the peak of A-A optimality, whereas the intervention may have no effect or be counterproductive for high-arousal SOS trials, which already surpass the optimal level, and thus additional arousal may be ineffective or disruptive.

To test these hypotheses, we leveraged real-time readouts of trial-by-trial pupil size to detect suboptimal and optimal A-A states and to trigger, on a random subset of SOS trials, a reorienting cue just prior to the retrieval attempt. After completing a goal-directed memory encoding task (Fig 1a), participants (*n* = 67) encountered studied and novel items and engaged in episodic retrieval. Specifically, on each retrieval trial, participants indicated whether they remembered the test probe as having been encountered in one of the two orienting tasks during encoding, they recognized the probe as having been studied but cannot remember the orienting task, or they perceived the probe as novel. Memory for studied items was assayed at the trial-level (hits/misses) and overall condition-level performance was computed using item memory *d*^*′*^ and source memory *d*^*′*^ (Fig 1b).

At the beginning of each retrieval block, we built participant-and-block-specific distributions of baseline-pupil during fixation periods, which we used to establish the trigger threshold for that block. Subsequently, we computed mean pupil size over a 1-s sampling window before each retrieval trial to detect, in real time, whether moment-to-moment pupillary dilation/constriction exceeded or fell within an empirical threshold, thus marking a trial as a SOS or OS event (Fig 1c). If the first window showed no SOS or had too much missing data, a second 1-s window followed. Either way, the window that determined the classification ended about 1.4 s before the retrieval probe, immediately before the trigger period. On 50% of SOS trials, PRISM triggered deployment of a salient, real-time arousal/attention-reorienting cue just prior to retrieval probe presentation. This design yielded three within-subjects conditions at retrieval: OS trials (optimal arousal/attention maintained throughout the real-time monitoring window; no intervention needed), SOS trials with trigger (suboptimal A-A state detected and arousal/attention reorienting cue delivered), and SOS trials without trigger (suboptimal A-A state detected but no reorienting cue delivered). Trigger and no-trigger SOS conditions were balanced by design via adaptive randomization that adjusts in real time to the history of past SOS trials. On each SOS trial, if the cumulative trigger/no-trigger distribution was at parity (50/50) for the block, assignment was determined by a random coin flip; the following SOS trial was then deterministically assigned to the underrepresented condition, restoring parity. Consequently, each successive pair of SOS trials contained exactly one trigger and one no-trigger trial in random order, which ensured that the cumulative distribution never deviated by more than one trial from parity. OS trials did not interact with the trigger assignment system and thus did not affect the trigger/no-trigger sequence.

The binary classification of trials as SOS or OS belies a potentially more nuanced reality; the inverted-U framework implies that A-A states vary continuously and that suboptimal states could differ in severity. That is, some represent moderate deviations from a peak A-A state and may reflect SOS states that remain within a recoverable range, while others represent extreme departures where intervention may be insufficient to restore performance. PRISM’s real-time detection algorithm, which applies a fixed threshold to tonic pupil size in real time, treats all threshold crossings equivalently. Theoretically, a trial that minimally crosses the SOS threshold may occupy a fundamentally different position on the A-A continuum than one representing a deep departure. Moreover, because tonic pupil size often exhibits slow non-stationary drift over the course of each block [70], some threshold crossings could be driven primarily by drift rather than putative A-A state changes.

During offline analysis, we further explored potentially time-varying A-A states to operationalize the severity continuum (see Fig 2). Because our real-time thresholds assumed a stationary A-A landscape defined at each block onset, slow time-on-task drift in tonic pupil size could by itself push trials outside the fixed thresholds as the block progressed. We therefore considered an alternative classification scheme: that the optimal A-A state itself shifts gradually over the course of a block. To estimate this moving distributional mean A-A level proxied by pupil size, we fit a spline to each participant’s pupil time series and reclassified each trial’s A-A state against the resulting drift-corrected thresholds (see **Materials and methods**). Each trial could thus be evaluated against two reference frames: the block-onset distributional mean (real-time, uncorrected) and the locally estimated distributional mean (drift-corrected), enabling us to distinguish between trials whose real-time SOS classification was driven by drift (i.e., representing putative moderately suboptimal states closer to the peak of the inverted-U) from those whose classification survived the correction (i.e., those representing putative deep suboptimal states further from the peak). This reclassification, when crossed with cue delivery, enabled post hoc testing of whether the reorienting cue’s functional impact scales with putative position on the A-A continuum, conferring greater benefit to moderately suboptimal states where the fixed-magnitude perturbation may be sufficient to nudge the state toward optimal performance, and lesser benefit to deeply suboptimal states where the same perturbation may be insufficient to bridge the larger deviation. Moreover, because the reclassification preserved the sign of the pupil deviation (constriction vs. dilation), it also enabled a direct test of any directional asymmetry in the effect of reorienting cues, which predicts that the A-A-boosting cue would selectively rescue low-A-A (constriction) SOS trials, which sit on the ascending limb of the inverted-U and can be nudged toward the peak, but not high-A-A (dilation) SOS trials, which sit on the descending limb where additional arousal/attention may be ineffective or counterproductive.

**Fig 2.**
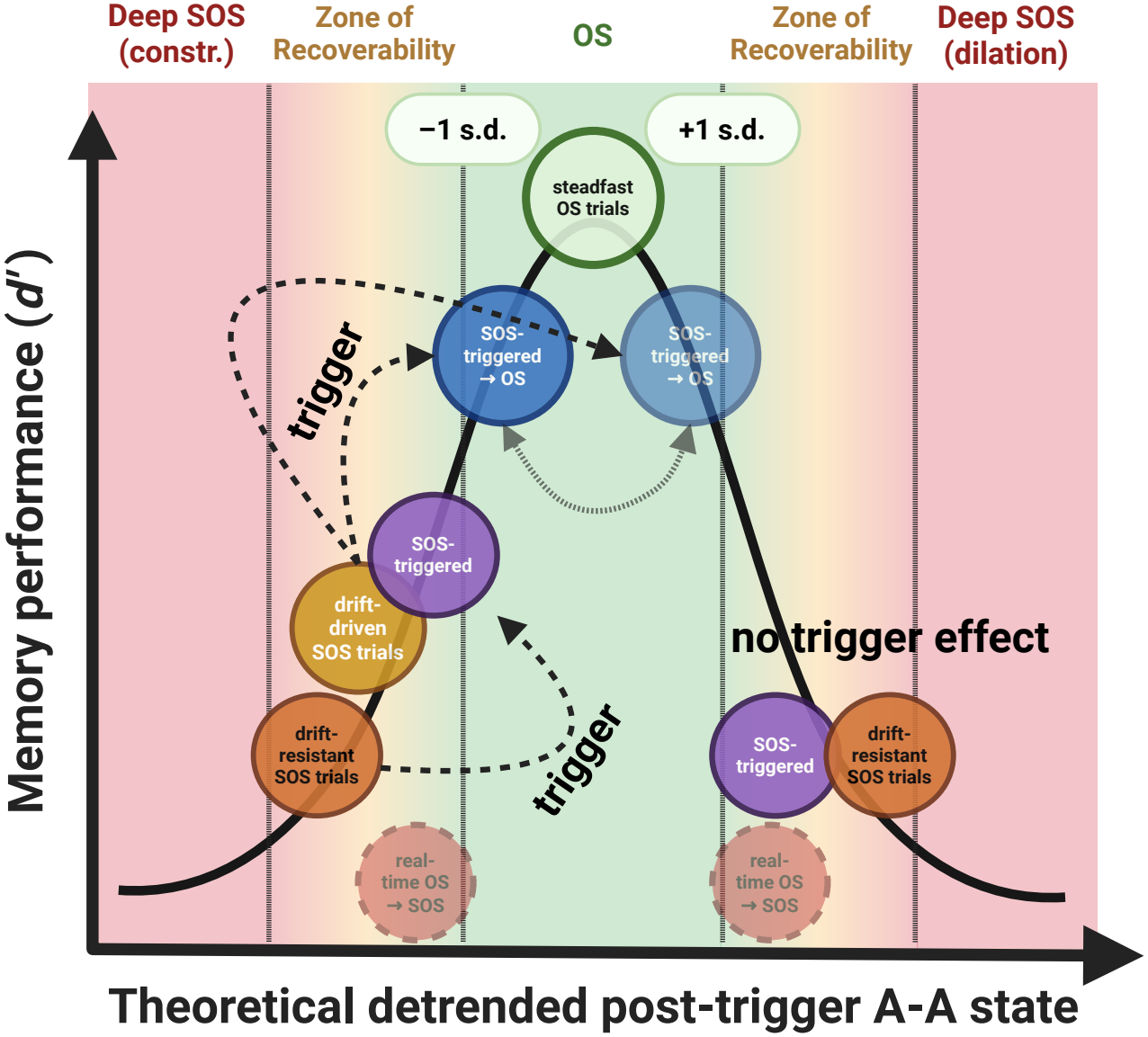
Theoretical framework mapping drift-corrected trial classifications onto an inverted-U shaped arousal-attentional (A-A) state continuum and their predicted memory performance. The x-axis represents the theoretical detrended post-trigger A-A state, ranging from deep suboptimal states (SOS) driven by pupil constriction (left) to deep SOS driven by pupil dilation (right), with the optimal state (OS) zone at center (bounded by *±*1 s.d.). The y-axis represents memory performance (item *d*^*′*^). The solid black curve depicts the predicted inverted-U relationship between detrended A-A state and memory performance. The six trial categories from the drift-corrected reclassification are positioned according to their inferred A-A states. Drift-resistant SOS trials (orange circles) occupy both deep SOS extremes, reflecting A-A departures severe enough to survive drift correction. Drift-driven SOS trials (yellow circle) fall within the zone of recoverability on the constriction side, indicating that their original SOS classification was attributable primarily to drift rather than a putative A-A state departure. SOS-triggered trials that remained SOS after drift correction (purple circles) appear at both extremes, whereas SOS-triggered trials reclassified as OS (blue circles) sit within or near the OS boundary on both sides. Steadfast OS trials (green circle) occupy the center of the OS zone. Real-time OS *→* SOS trials (faded red circles) are depicted off the inverted-U curve to reflect that their position along the A-A state continuum does not conform to the predicted mapping. Although these trials exhibited the lowest *d*^*′*^ of any condition, their mean detrended pupil deviations were closer to zero than those of the drift-resistant SOS trials, placing them nearer to the OS zone rather than at the deep SOS extremes. This discrepancy likely arises from a selection bias inherent to spline-based reclassification: OS *→* SOS reclassification requires that the spline trend shifted away from the real-time baseline, and trials on the same side as this shift are mechanically constrained to be closer to the corrected mean than trials whose optimality status survived correction. The reduced opacity and placement below the curve further reflect the instability of *d*^*′*^ estimates for this condition due to low trial counts (527 total trials across all participants). Dashed arrows illustrate the effect of the reorienting cue, which elevates performance for trials within the zones of recoverability but has no effect on trials in the deep SOS dilatory extreme, where the A-A state departure is too severe for a brief visual reorienting cue to remediate.

The closed-loop design tests whether perturbing A-A state changes memory. We complemented it with two correlational analyses that describe how memory varies with A-A state within and across participants. First, on triggered trials, we asked whether pupil size during the 1 s between the reorienting cue and the retrieval probe predicted whether a studied item was remembered. This pre-probe pupil indexes the A-A state in which each retrieval attempt began, after the cue had acted. If the cue raises arousal, the inverted-U framework implies that it could leave some SOS trials short of the optimum and push others past it. Memory on triggered trials should then vary with pre-probe pupil size, and be poorest on trials the cue pushed furthest past the optimum. Second, we asked whether trait-level differences in the tendency for attention to lapse explain, in part, why some individuals remember more than others. Readiness-to-remember accounts address variability in memory across individuals as well as across moments [5], and prior work linked individual differences in sustained attention and media multitasking to memory failure [6]. To measure trait attention, participants completed two sustained attention tasks and questionnaires on media multitasking, attention-deficit/hyperactivity disorder (ADHD) symptoms, impulsivity, attentional control, and mind wandering after the memory task, and we combined these measures into a single latent attention factor.

## Results

### Real-time closed-loop A-A state intervention at retrieval

First, we computed item and source memory sensitivity (*d*^*′*^; log-linear corrected) for each participant in each of the three real-time retrieval conditions (OS, SOS without trigger, SOS with trigger) and fit a Bayesian linear model with a by-participant random intercept to estimate each condition’s mean and test directional hypotheses (see **Materials and methods**). For each condition we report the posterior median (*m*) and its 95% highest-density interval (HDI). For each contrast we report the posterior mean difference (*b*), its 90% credible interval (CrI), and its evidence ratio (ER), the posterior probability of the predicted direction divided by that of the opposite direction. We call a contrast credible when its 90% CrI lies entirely on the predicted side of zero, and we describe ERs of 1–3, 3–10, 10–30, 30–100, and above 100 as anecdotal, moderate, strong, very strong, and extreme evidence. Item memory *d*^*′*^ was credibly higher on OS trials (*m* = 2.64, 95% HDI [2.46, 2.81]) than on SOS trials without trigger (*m* = 2.39, 95% HDI [2.22, 2.57]; *b* = *−*0.25, 90% CrI [*−*0.37, *−*0.12], ER *>* 1,000), providing extreme evidence for an effect of attentional state on memory performance (Fig 3a–c). Source memory *d*^*′*^ was also credibly lower on SOS trials without trigger (OS: *m* = 2.14, 95% HDI [1.97, 2.31]; SOS without trigger: *m* = 2.02, 95% HDI [1.85, 2.19]; *b* = *−* 0.12, 90% CrI [*−*0.24, *−*0.01], ER = 23.54), providing strong evidence that preparatory lapsing at retrieval impaired not only item memory but also recollection of the encoding context (Fig 3d–f).

**Fig 3.**
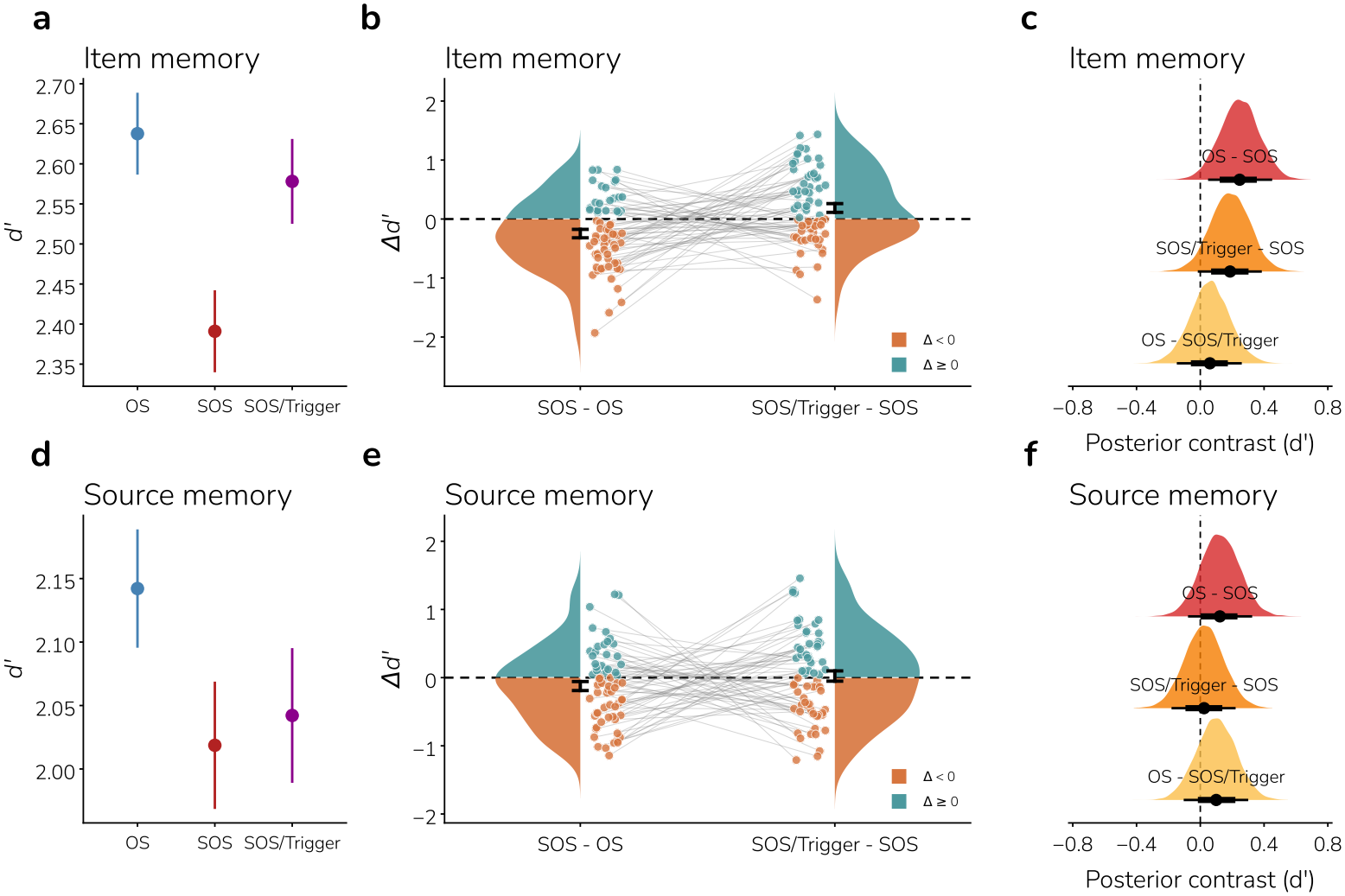
Real-time reorienting cues recover item memory but not source memory from pupil-defined arousal-attentional (A-A) suboptimal states (SOS). SOS and trigger recovery effects for item memory (**a**,**b**) and source memory (**d**,**e**) *d*^*′*^, and estimated marginal contrasts for *d*^*′*^ (**c**,**f**) between the three within-subjects conditions at retrieval. Plots of estimated marginal contrasts show posterior probability distributions. Points indicate posterior medians; thick and thin horizontal lines indicate the 66% and 90% HDIs, respectively. Dashed vertical lines mark zero (no difference).

Next, we tested whether SOS trials with triggers showed better memory than SOS trials without triggers, the central causal test of whether our pre-retrieval reorienting cue rescues memory access in real time. Item memory *d*^*′*^ was credibly higher on SOS trials with triggers (*m* = 2.58, 95% HDI [2.41, 2.76]) than on SOS trials without triggers (*m* = 2.39, 95% HDI [2.22, 2.57]; *b* = 0.19, 90% CrI [0.06, 0.31], ER = 163.38), providing extreme evidence that delivering an attentional reorienting trigger before the retrieval attempt improved memory for items that would otherwise be impaired by a suboptimal A-A state. Item memory on triggered SOS trials approached that on OS trials (*m* = 2.64, 95% HDI [2.46, 2.81]; *b* = *−*0.06, 90% CrI [*−*0.18, 0.06], ER = 3.99). However, contrary to our hypothesis, the triggers did not credibly rescue source recollection (SOS with trigger: *m* = 2.04, 95% HDI [1.88, 2.22]; SOS without trigger: *m* = 2.02, 95% HDI [1.85, 2.19]; *b* = 0.02, 90% CrI [*−* 0.09, 0.14], ER = 1.70), and the evidence for any benefit was anecdotal. Source *d*^*′*^ on triggered SOS trials also remained below that on OS trials (*b* = *−* 0.10, 90% CrI [*−* 0.22, 0.01], ER = 12.16). Moreover, the trigger effect was credibly larger for item than for source memory (item minus source trigger effect: *b* = 0.16, 90% CrI [0.06, 0.26], ER = 249.00), providing extreme evidence that the cue benefited item memory more than source memory.

### Offline drift correction reclassifies real-time trials

A fundamental challenge in closed-loop physiological monitoring is the potential for systematic shifts that impact real-time classification algorithms; specifically, pupil measures are susceptible to slow physiological drift (i.e., the “time-on-task effect”) [51, 71–77]. Our primary analysis relied on a causal, real-time pupillometry-based algorithm to detect A-A SOS and deliver visual reorienting cues on a subset of SOS trials. The real-time algorithm used a fixed baseline computed from the first 20 “burn-in” trials of each retrieval block to identify subsequent moments when pupil size fell above/below 1 s.d. around the baseline mean. Given that drift can arise from factors such as fatigue, autonomic regulation, and luminance adaptation, to name a few, it is not unreasonable to suspect that within-participant slow trends could impact real-time SOS detection in structured ways. Critically, the use of a fixed early-block baseline (rather than an adaptive rolling baseline, which has its own challenges) would be impacted by drifts such that trials later in the block would be increasingly likely to fall outside the fixed threshold and therefore be classified as SOS trials, even when the participant’s A-A state relative to their time-varying physiological baseline has not changed. This design choice, while necessary for real-time implementation in the present study, warrants rigorous post hoc examination.

To validate the robustness of PRISM’s SOS detection and to operationalize a continuum of suboptimality, we implemented an offline spline drift correction that reclassified trials independent of slow pupil drift. For each participant and block, we constructed a continuous time series by concatenating the epoched pupil data from the real-time monitoring windows across all trials in trial order, with inter-trial gaps padded with NaN values to preserve the temporal structure of the block (padding duration: 10,416 ms per trial, corresponding to the trigger window, post-trigger fixation, stimulus, and pre-stimulus intertrial intervals). A natural cubic spline (*df* = 5) was then fit to this padded time series using only the non-NaN samples from the real-time monitoring windows, capturing slow time-varying trends including linear drift and exponential decay components (Fig 4a).

**Fig 4.**
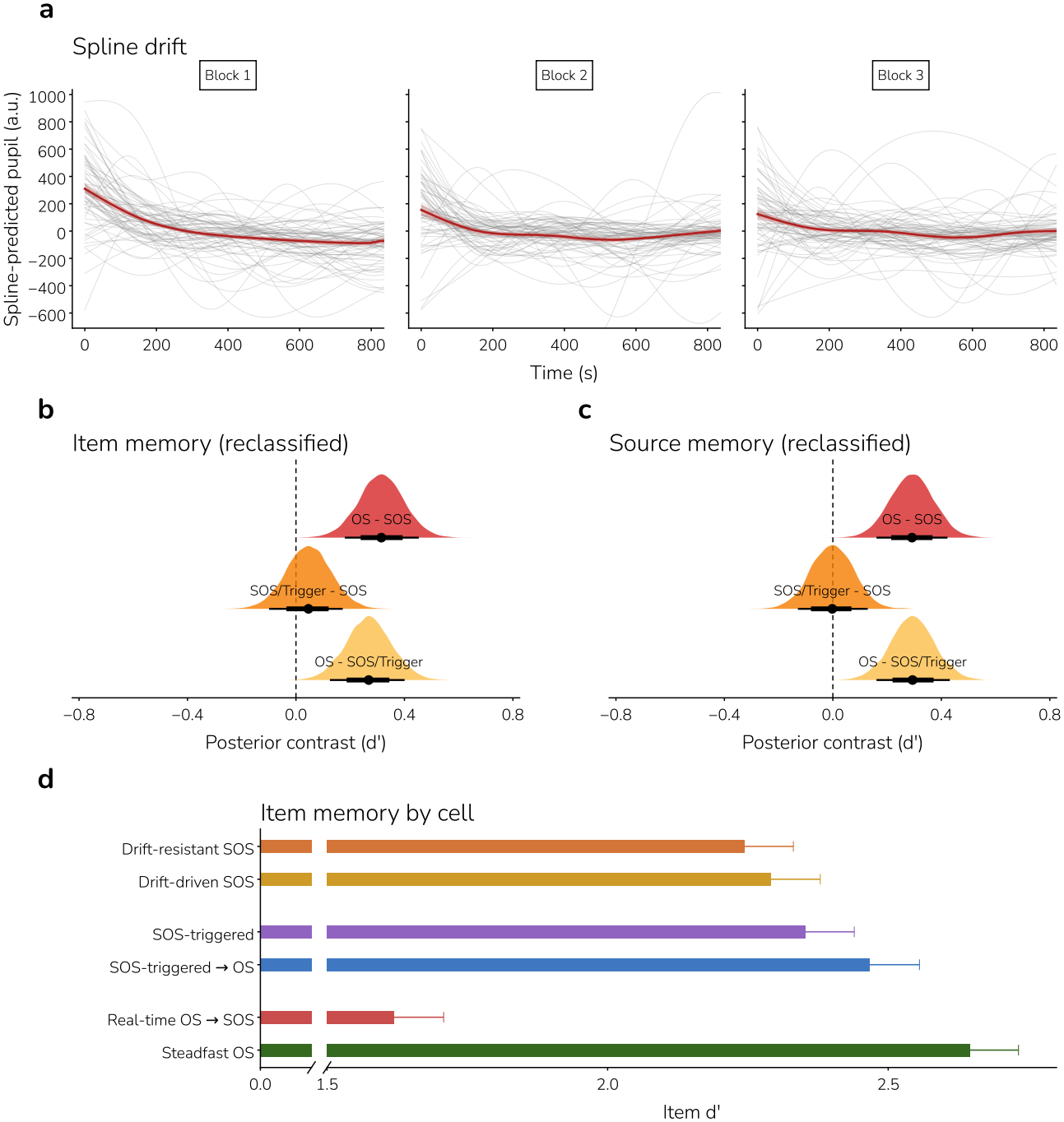
Pupil drift characteristics and memory performance after drift-corrected reclassification of arousal-attentional states. (**a**) Mean-centered spline-predicted pupil size (arbitrary units) across each of the three experimental blocks. Grey lines depict individual participants; the red line and ribbon denote the grand mean *±*1 s.e.m. Spline fits capture slow, non-stationary drift in the pupil signal that the real-time classification algorithm did not account for. The x-axis is trimmed to 800 s to represent the stable portion of the drift grand mean (as there were varying block lengths per participant — with few spanning beyond 800 s — depending on the number of “recycled” trials one encounters in the real-time paradigm). (**b**,**c**) Posterior distributions of estimated marginal contrasts for item memory (**b**) and source memory (**c**) *d*^*′*^ between the three within-subjects conditions after drift-corrected reclassification (optimal state, suboptimal state, suboptimal state with trigger). Points indicate posterior medians; thick and thin horizontal lines indicate the 66% and 90% HDIs, respectively. Dashed vertical lines mark zero (no difference). (**d**) Posterior mean item *d*^*′*^ for each of the six cells of the real-time/reclassified transition table (Table 1), from a Bayesian cell-means model fit to participant-level item *d*^*′*^ (log-linear corrected). Error bars denote *±*1 posterior s.d.; the *x*-axis is broken between 0 and 1.5. Cells are grouped by real-time classification and listed in the order of Table 1. In each real-time SOS group, posterior mean *d*^*′*^ is lower for the cell that remained SOS after drift correction (drift-resistant) than for the cell reclassified as OS (drift-driven). Steadfast OS trials showed the highest posterior mean *d*^*′*^ of any cell and real-time OS *→* SOS trials the lowest, though this may partly reflect selection artifacts of the spline-based reclassification (see Fig 2 caption).

For each trial, we computed the mean spline prediction during that trial’s real-time monitoring window, representing the estimated drift level at the time of that trial. Each trial’s monitoring-window pupil mean was then corrected by subtracting the trial-specific spline mean. Unlike the real-time algorithm, which computed thresholds from a baseline initialized over the first 20 burn-in trials, the reclassification thresholds were derived from the global distribution of spline-detrended pupil values across all real-time monitoring samples within the block: trials were classified as SOS if the drift-corrected monitoring-window mean fell above/below 1 s.d. of the global detrended mean. This procedure leverages information from the entire block — including trials that were unavailable to the real-time classifier — providing an alternate drift-corrected reference for threshold computation.

**Table 1.** Six-cell condition matrix.

| **Original ↓ Reclassified →** | **SOS** | **SOS-triggered** | **OS** | **OS-triggered** |
|---|---|---|---|---|
| **SOS** | 2,032 | 0 | 2,445 | 0 |
| **SOS-triggered** | 0 | 2,037 | 0 | 2,430 |
| **OS** | 527 | 0 | 3,610 | 0 |

Crossing the real-time and drift-corrected classifications of trials produced a six-cell condition matrix (Table 1). Of the 4,477 real-time SOS-untriggered trials, 2,032 (45.4%) retained their SOS classification after drift correction. We term these *drift-resistant SOS-untriggered* trials. Their A-A state departures were extreme enough that removing the drift component did not change their classification. The remaining 2,445 (54.6%) were reclassified as OS. We term these *drift-driven SOS-untriggered* trials, because their original SOS classification reflected mainly drift rather than a substantial A-A state departure. Among the 4,467 real-time SOS-triggered trials, 2,037 (45.6%) remained SOS (*drift-resistant SOS-triggered* trials) and 2,430 (54.4%) were reclassified as OS (*drift-driven SOS-triggered* trials). Of the 4,137 real-time OS trials, only 527 (12.7%) were reclassified as SOS after drift correction. In these trials, drift had shifted the raw pupil trace toward the baseline mean and masked a suboptimal A-A state that appeared once the drift was removed. The other 3,610 (87.3%) retained their OS classification (*steadfast OS* trials).

We then examined whether the SOS effects on item and source memory *d*^*′*^ could be replicated with the drift-corrected trial reclassification (Fig 4b,c). Under reclassification, there was extreme evidence that item memory *d*^*′*^ was lower on SOS than on OS trials (OS: *m* = 2.62, 95% HDI [2.45, 2.79]; SOS without trigger: *m* = 2.30, 95% HDI [2.13, 2.48]; *b* = *−*0.31, 90% CrI [*−*0.45, *−*0.18], ER *>* 1,000; Fig 4b). Source memory *d*^*′*^ also showed an SOS impairment with extreme evidence (OS: *m* = 2.19, 95% HDI [2.02, 2.36]; SOS without trigger: *m* = 1.90, 95% HDI [1.73, 2.07]; *b* = *−* 0.29, 90% CrI [*−*0.42, *−* 0.16], ER *>* 1,000), more than twice the size of the source effect under the real-time classification (*b* = 0.12; Fig 4c).

Next, we examined whether the trigger benefit for item memory in the real-time analysis replicated with the drift-corrected classification. Under reclassification, the evidence for a trigger effect on item memory was anecdotal (SOS without trigger: *m* = 2.30, 95% HDI [2.13, 2.48]; SOS with trigger: *m* = 2.35, 95% HDI [2.18, 2.52]; *b* = 0.05, 90% CrI [*−* 0.09, 0.18], ER = 2.49) using all SOS trials including the OS trials reclassified as SOS. ^1^ Source memory continued to show no trigger benefit as for real-time SOS trials before (SOS without trigger: *m* = 1.90, 95% HDI [1.73, 2.07]; SOS with trigger: *m* = 1.90, 95% HDI [1.73, 2.06]; *b* = 0.00, 90% CrI [*−* 0.13, 0.12], ER = 0.96), and with an ER near 1 the data favored neither direction.

Unlike the weak evidence in the SOS trials under reclassification, a clear trigger benefit was observed in the drift-driven SOS trials, which the real-time algorithm flagged as SOS but drift correction reclassified as OS. On these trials, the trigger improved item *d*^*′*^ with strong evidence (ER = 29.85; see below). Despite the reduced number of trials for the comparisons under reclassification, this unexpected finding of the previously credible benefit of delivering a reorienting cue in real time on SOS trials splitting into two qualitatively different trigger effects invites further investigation. We wondered whether this offline reclassification of A-A states is revealing a more complex underlying structure of the real-time trigger effect on item memory, which motivated the follow-up analyses in the next section.

### Item memory across the six reclassified cells

If the drift-corrected classifications capture a meaningful continuum of A-A state quality, then item memory *d*^*′*^ should vary systematically across the six classification cells. Cell membership was set by pupil size alone. The real-time and drift-corrected thresholds were applied to the pupil in the sampling window before any cue, so each SOS trial’s depth label (drift-resistant or drift-driven) was assigned without reference to memory performance. These labels describe the A-A state before the cue. Because the cue then changes that state, we do not infer how far a triggered trial was from optimality at retrieval, and we compare depths only within a trigger condition. We computed item memory *d*^*′*^ (log-linear corrected) for each participant in each cell and fit the same Bayesian model, with a by-participant random intercept, to estimate each cell’s mean and test directional hypotheses (see **Materials and methods**).

Posterior mean item *d*^*′*^ was lower for drift-resistant than for drift-driven SOS trials within each trigger condition, and higher for triggered than for untriggered trials within both drift-resistant and drift-driven SOS (Fig 4d; contrasts below). We interpret the drift-resistant versus drift-driven differences as reflecting different depths of suboptimality before the cue. Steadfast OS trials showed the highest *d*^*′*^ of any condition (*m* = 2.65, 95% HDI [2.47, 2.81]). Real-time OS trials reclassified as SOS showed the lowest (*m* = 1.62, 95% HDI [1.45, 1.80]).

For drift-driven SOS trials, which likely fall closer to A-A state optimality than drift-resistant SOS trials, the trigger credibly improved item *d*^*′*^, with strong evidence (triggered vs. untriggered: *b* = 0.18, 90% CrI [0.02, 0.33], ER = 29.85). For drift-resistant SOS trials, which likely fall further from optimality, the evidence for a trigger benefit was moderate and in the same direction (triggered vs. untriggered: *b* = 0.11, 90% CrI [*−*0.05, 0.26], ER = 7.03), although the 90% CrI included zero. The interaction contrasting these two trigger effects was inconclusive (*b* = 0.07, 90% CrI [*−*0.15, 0.28], ER = 2.27), leaving open whether the trigger operates differently at different depths of suboptimality. A fixed-perturbation account holds that a brief A-A-boosting stimulus produces greater functional recovery when the distance to the inverted-U peak is smaller. It therefore predicts a larger trigger benefit for drift-driven than for drift-resistant SOS states. The present data are directionally compatible with this prediction but lack the statistical evidence to adjudicate it.

Drift-resistant SOS trials showed lower item *d*^*′*^ than drift-driven SOS trials both without the trigger (*b* = *−* 0.05, 90% CrI [*−* 0.20, 0.10], ER = 2.32) and with it (*b* = *−* 0.12, 90% CrI [*−* 0.27, 0.04], ER = 8.27), with anecdotal and moderate evidence, respectively, although neither difference was credible. The difference was larger among triggered trials because the trigger raised item *d*^*′*^ more for drift-driven than for drift-resistant trials, although the interaction testing this was inconclusive (see above). Comparisons with steadfast OS trials were clearer. Steadfast OS trials showed credibly higher *d*^*′*^ than both drift-driven SOS-triggered trials, with very strong evidence (*b* = 0.18, 90% CrI [0.02, 0.33], ER = 35.92), and drift-resistant SOS-triggered trials, with extreme evidence (*b* = 0.29, 90% CrI [0.14, 0.45], ER *>* 1,000). Even with the trigger, neither drift-driven nor drift-resistant SOS trials recovered to the level of steadfast OS trials.

### Asymmetric effects across hypo vs. hyperarousal suboptimal states

The inverted-U framework predicts that the trigger’s efficacy should depend not only on the magnitude of suboptimality but also on its direction. Hypoarousal suboptimal states (pupil constriction), which sit on the ascending limb of the inverted-U, should be more amenable to an A-A-boosting intervention than hyperarousal suboptimal states (pupil dilation), which sit on the descending limb where additional arousal might be counterproductive. We tested this planned hypothesis across all real-time SOS trials, examining the trigger’s effect as a function of whether a trial’s SOS classification was due to pupil constriction or pupil dilation. The trigger credibly improved *d*^*′*^ for constriction trials, with strong evidence (*b* = 0.18, 90% CrI [0.002, 0.37], ER = 19.51), whereas the evidence for a dilation boost was anecdotal (*b* = 0.05, 90% CrI [*−* 0.13, 0.24], ER = 2.20). The interaction was in the predicted direction, with moderate evidence for a larger trigger boost for constrictions than dilations (*b* = 0.13, 90% CrI [*−* 0.13, 0.39], ER = 3.92), although its 90% CrI included zero. Post hoc, we split this test by depth of suboptimality. Among drift-resistant SOS trials (i.e., the most deeply suboptimal states), the same pattern appeared: a credible constriction boost with strong evidence (*b* = 0.19, 90% CrI [0.01, 0.37], ER = 24.53), no credible dilation boost (*b* = 0.06, 90% CrI [*−* 0.13, 0.24], ER = 2.24), and moderate evidence for the interaction (*b* = 0.14, 90% CrI [*−* 0.12, 0.40], ER = 4.27). This pattern is consistent with the prediction that suboptimality is not sign-neutral with respect to recovery. A brief A-A-boosting cue may nudge low-A-A states toward the inverted-U peak, whereas the same perturbation is less effective, or potentially counterproductive, for states already at the high-A-A extreme. Because neither interaction was credible, however, the asymmetry itself remains suggestive. Among drift-driven SOS trials (i.e., the marginally suboptimal states nearer to the peak), the evidence was anecdotal throughout (constriction: *b* = 0.09, 90% CrI [*−* 0.15, 0.33], ER = 2.69; dilation: *b* = 0.00, 90% CrI [*−* 0.25, 0.26], ER = 1.03; interaction: *b* = 0.08, 90% CrI [0.26, 0.43], ER = 1.92). These data therefore cannot tell whether the direction of a modest departure from optimality matters, although the trigger’s overall benefit for drift-driven trials was credible (ER = 29.85; see above). An interesting open question is whether the cue works by increasing both arousal and attention in the hypo-A-A arm but only attention, without adverse further increases in arousal, in the hyper-A-A arm.

### Pre-probe pupil size and trial-level memory outcomes

In order to assess whether the magnitude of pupil size in response to the reorienting cue further explains memory performance across the intervened trials, we asked whether pupil size immediately before the retrieval probe predicted memory on individual real-time SOS trials with triggers. For each retained trial, we averaged the preprocessed, block-wise *z*-scored pupil over the 1-s post-trigger fixation that preceded the object probe (hereafter, pre-probe pupil; see **Materials and methods**). There was extreme evidence that larger pre-probe pupil predicted a lower probability of remembering (*b* = *−*0.17, 90% HDI [*−*0.26, *−*0.07], ER = 427.57; Fig 5a–d). Item memory primarily drove this relationship (item slope: *b* = *−*0.23, 90% HDI [*−*0.35, *−*0.11], ER = 799.00; source slope: *b* = *−*0.10, 90% HDI [*−*0.21, *−*0.003], ER = 18.93; pupil *×* memory: *b* = 0.12, 90% HDI [*−*0.01, 0.26], ER = 13.71), and it held in a separate item-only model (*b* = *−*0.22, 90% HDI [*−*0.36, *−*0.09], ER = 180.82). Thus, among real-time triggered SOS trials, pre-probe A-A states with the largest pupils were the least conducive to item memory. These are plausibly the states that the A-A-boosting reorienting cue pushed further toward the high-arousal-attentional extreme, consistent with the asymmetric trigger effects reported above.

**Fig 5.**
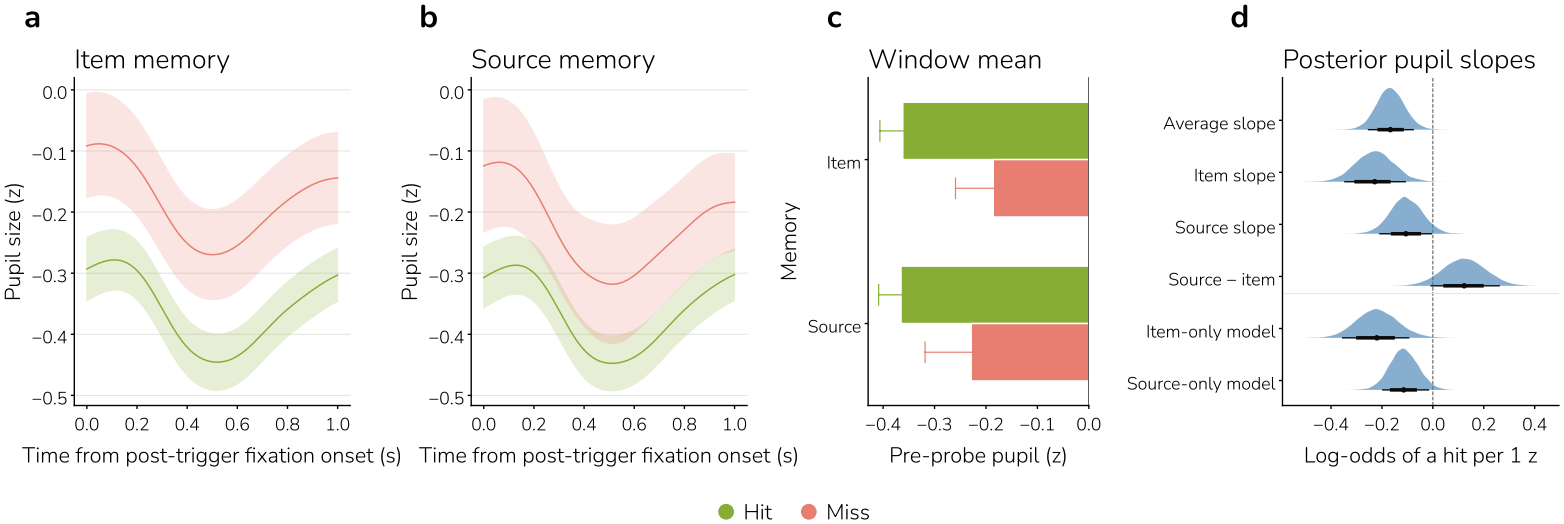
Pre-probe pupil size and memory on SOS-triggered trials. Pupil size during the 1-s fixation between the reorienting cue and the retrieval probe, on trials where the real-time system detected a SOS and delivered the cue. Only studied items with no missing pupil samples in the window are included (2,240 trials from 67 participants). (**a, b**) *z*-scored pupil size over the fixation for item (**a**) and source (**b**) hits and misses. Lines and bands show the mean *±* 1 s.e.m. across participants of each participant’s average trace. (**c**) Mean pupil size over the window, which is the predictor in **d**. It is averaged within participants, then shown as the mean *±* 1 s.e.m. across participants. (**d**) Posterior pupil slopes from Bayesian logistic mixed models of hit vs. miss, in log-odds of a hit per 1-s.d. increase in pupil size. Above the line are terms from the pupil *×* memory type model: the average slope, the item and source slopes, and their difference. Below the line are separate models fit to item and source trials alone. Shapes are posterior densities, points are medians, and thick and thin bars are the 66% and 90% HDIs.

### A latent attention factor predicts item memory

The analyses above concern fluctuations in A-A state within each participant. We next asked a between-participant question: do people with better trait attention remember more? Prior work linked individual differences in sustained attention and media multitasking to memory failure [6], and we adapted that study’s individual-differences assessment, which participants completed after the memory task (see **Materials and methods**). It comprised two sustained attention tasks and a set of questionnaires. In the gradual-onset continuous performance task (gradCPT) [39], participants watched a stream of gradually blending scenes, pressing a key for frequent city scenes and withholding responses to rare mountain scenes. In the sustained attention to response task (SART) [78], participants classified a rapid stream of shape displays in which one shape type was rare. The questionnaires measured media multitasking, ADHD symptoms, attentional impulsivity, everyday attentional control, and mind wandering.

Each of these measures is a noisy and partial index of attention, and testing every one against both memory outcomes would require 20 separate comparisons. We therefore combined them with a Bayesian one-factor confirmatory factor analysis (Fig 6a). The model treats each measure as an indicator, an observed score assumed to reflect a single unobserved trait, and estimates that trait from the variance the indicators share. Each indicator’s loading describes how strongly it reflects the trait. We used ten indicators, one per construct. Where a task or questionnaire yielded several overlapping scores, such as gradCPT *d*^*′*^ and the error rates it is computed from, we kept one, so that no construct entered the model twice (see **Materials and methods**). The three task-based indicators were gradCPT sensitivity (*d*^*′*^), gradCPT response-time variability (RT CoV) [39], and SART rare-shape accuracy [78]. The seven self-reported indicators were media multitasking (Media Multitasking Inventory) [79], ADHD symptoms [80], attentional impulsivity (Barratt Impulsiveness Scale-11, BIS-11) [81], attentional control over distraction and shifting [82], and spontaneous and deliberate mind wandering [82]. Each indicator was signed so that higher values denote better attention.

**Fig 6.**
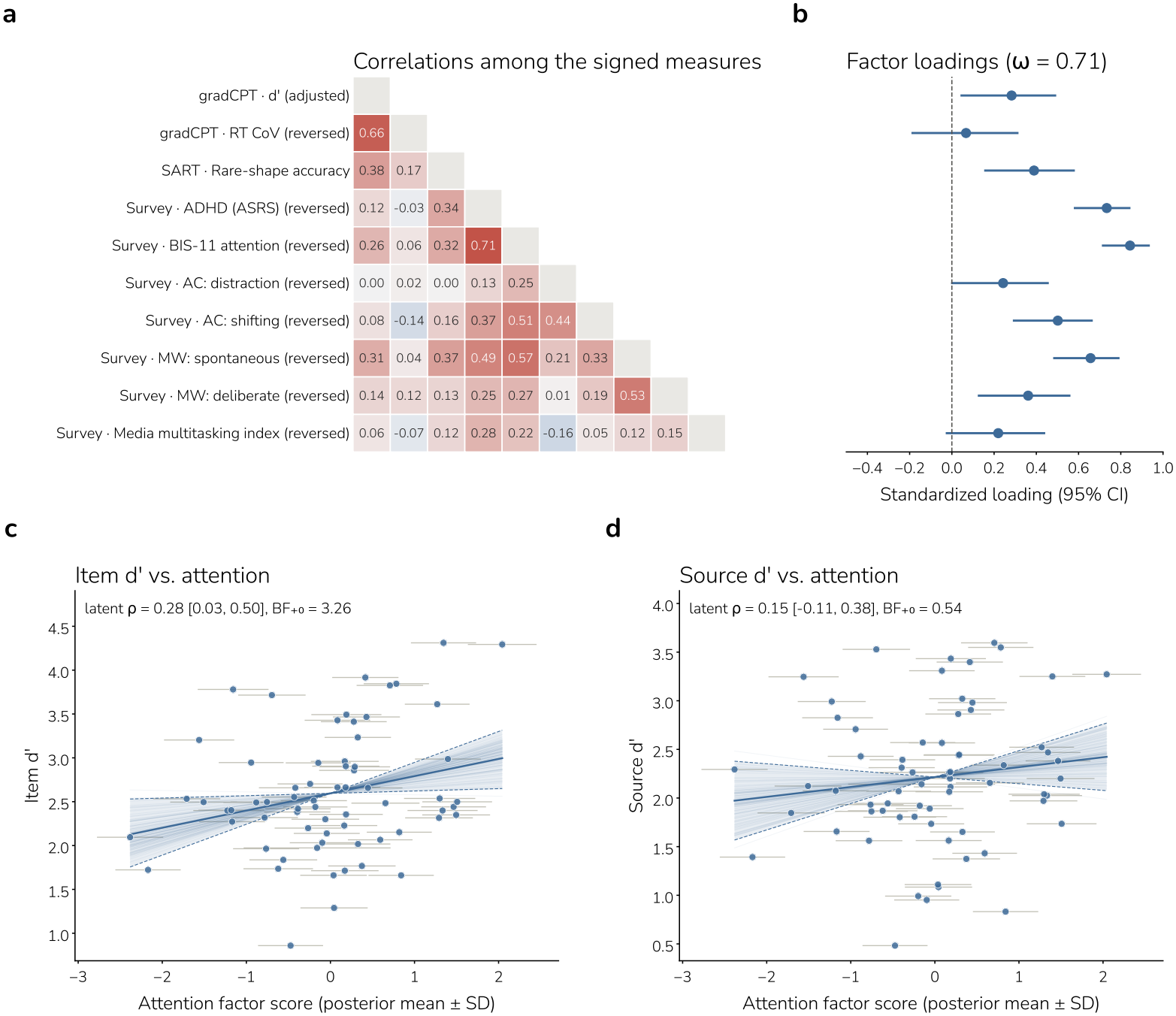
A latent attention factor and memory. A Bayesian one-factor model was fit to ten measures of attention: three tasks (gradCPT *d*^*′*^ and RT CoV, SART rare-shape accuracy) and seven self-report surveys. Each measure was *z*-scored and signed so that higher means better attention. The model uses every observed score (*N* = 67; 60 completed all ten). (**a**) Pearson correlations among the signed measures (red, positive; blue, negative). AC, attentional control; MW, mind wandering. (**b**) Standardized loadings of each measure on the factor (posterior means with 95% credible intervals), in the same row order as **a**. *ω*, McDonald’s omega reliability of the factor (95% CrI [0.62, 0.78]). (**c, d**) Item (**c**) and source (**d**) *d*^*′*^ against each participant’s attention factor score. Points are posterior means and horizontal bars are *±* 1 posterior s.d. Latent *ρ* is the posterior mean correlation between the factor and *d*^*′*^ with its 95% credible interval. Faint lines are 100 posterior draws and the band is the 95% HDI. Because *d*^*′*^ enters the model *z*-scored, the fit passes through mean *d*^*′*^ at the mean factor score, so the band reflects uncertainty in the slope only. BF_+0_ is the directional Bayes factor for *ρ >* 0 (better attention, better memory) against *ρ* = 0.

All posterior mean loadings were positive, and their 95% credible intervals excluded zero for seven of the ten indicators (Fig 6b). They were largest for BIS-11 attention (*λ* = 0.84, 95% CrI [0.71, 0.94]), ADHD symptoms (*λ* = 0.73, [0.58, 0.85]), and spontaneous mind wandering (*λ* = 0.66, [0.48, 0.79]). Loadings were intermediate for attentional shifting (*λ* = 0.50, [0.29, 0.67]), SART rare-shape accuracy (*λ* = 0.39, [0.15, 0.58]), deliberate mind wandering (*λ* = 0.36, [0.12, 0.56]), gradCPT *d*^*′*^ (*λ* = 0.28, [0.04, 0.49]), attentional distractibility (*λ* = 0.24, [*−* 0.004, 0.46]), and media multitasking (*λ* = 0.22, [*−* 0.03, 0.44]). The gradCPT RT CoV contributed little (*λ* = 0.07, [*−* 0.19, 0.32]). Composite reliability was acceptable (*ω* = 0.71, 95% CrI [0.62, 0.78]), and Bayesian sampling diagnostics were satisfactory (all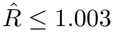, bulk ESS *>* 1,400, no divergent transitions). Higher factor scores tracked every raw indicator in the expected direction. Self-report dominated the factor (*r* = *−* 0.93 with BIS-11 attention and *r* = *−* 0.82 with ADHD symptoms), with smaller contributions from SART (*r* = 0.46) and gradCPT (*r* = 0.33) performance.

The attention factor predicted item memory *d*^*′*^ (standardized slope *β* = 0.27, 95% HDI [0.04, 0.52], BF_+0_ = 3.21; Fig 6c). For scale, participants in the top third of factor scores (*n* = 22) had a mean item *d*^*′*^ of 2.86, compared with 2.41 in the bottom third (*n* = 23), a difference of 0.44. A latent correlation estimated within the same structural model, which accounts for uncertainty in the factor scores, gave the same result (*ρ* = 0.28, 95% CrI [0.03, 0.50], BF_+0_ = 3.26), as did a unit-weighted composite of the same ten signed indicators (*β* = 0.34, 95% HDI [0.11, 0.58], BF_+0_ = 15.54).

Associations with source memory *d*^*′*^ were not credible (*β* = 0.14, 95% HDI [*−*0.11, 0.39], BF_+0_ = 0.50; latent *ρ* = 0.15, 95% CrI [*−* 0.11, 0.38], BF_+0_ = 0.54; Fig 6d). Neither a task-only nor a questionnaire-only factor yielded a credible latent correlation with item memory *d*^*′*^ (tasks: *ρ* = 0.22, 95% CrI [*−*0.04, 0.46], BF_+0_ = 1.27; questionnaires: *ρ* = 0.24, 95% CrI [*−* 0.02, 0.46], BF_+0_ = 1.96). The three estimates were similar (*ρ* = 0.22–0.28), and only the combined factor’s interval excluded zero, consistent with both modalities indexing the same memory-relevant attention trait.

To see which indicators carried the association with item memory, we conducted an exploratory analysis relating item and source memory *d*^*′*^ to each of the ten indicators separately (Fig 7). All 20 slopes pointed in the predicted direction, with better attention tracking better memory. SART rare-shape accuracy showed the strongest relation, with very strong evidence for item memory (*β* = 0.40, 95% HDI [0.18, 0.63], BF_+0_ = 79.24) and moderate evidence for source memory (*β* = 0.31, 95% HDI [0.08, 0.55], BF_+0_ = 7.60). Spontaneous mind wandering showed moderate evidence for item memory (*β* = *−*0.31, 95% HDI [*−*0.55, *−*0.08], BF_*−*0_ = 6.69) but not for source memory (BF_*−*0_ = 0.63). Evidence linking gradCPT *d*^*′*^ (BF_+0_ = 1.54) and BIS-11 attention (BF_*−*0_ = 1.11) to item memory was anecdotal, and the remaining 14 slopes had Bayes factors below 1 (0.17–0.70). The factor’s association with item memory therefore pools many small relations that all point the same way, with SART performance the largest single contributor. SART was also the only indicator related to source memory, and the factor, dominated by self-report, did not carry this relation.

**Fig 7.**
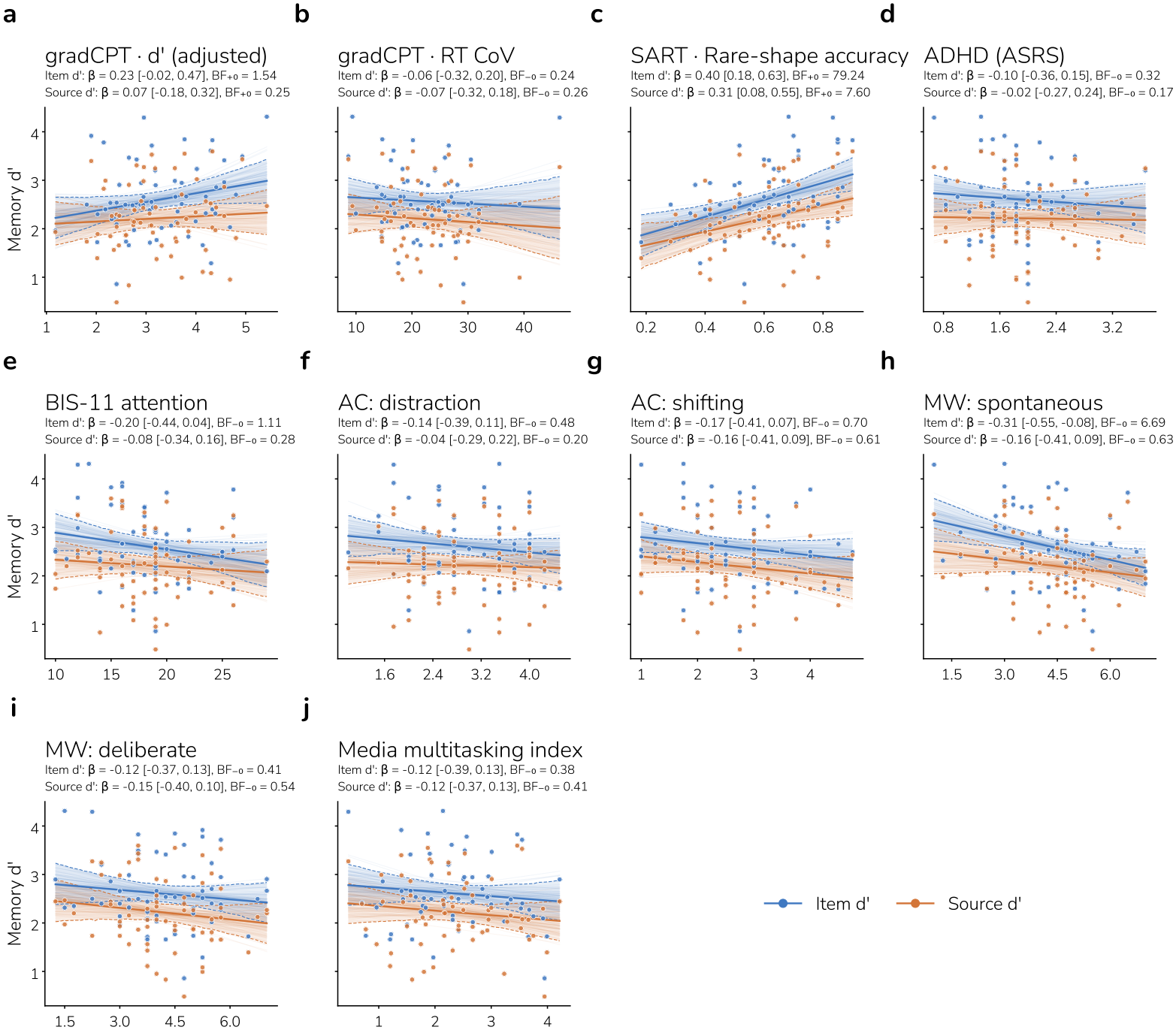
Memory and each measure in the attention factor. Item (blue) and source (orange) *d*^*′*^ against each of the ten measures that make up the latent attention factor in Fig 6, each on its raw scale (*n* = 63–66 participants per measure). For RT CoV and the seven surveys, higher scores mean worse attention, so the predicted relation (better attention, better memory) is a negative slope. Each line is a Bayesian linear regression of *d*^*′*^ on the measure (brms default priors [83]) at the posterior mean. Faint lines are 100 posterior draws, and dashed lines bound the 95% HDI. *β* is the standardized slope (posterior mean with 95% HDI). BF_+0_ (gradCPT *d*^*′*^ and SART accuracy) and BF_*−*0_ (all other measures) are one-sided Bayes factors for a correlation in the predicted direction against no correlation. They use a uniform prior on *ρ* over the predicted half and are computed from the Pearson correlation. (**a, b**) gradCPT. (**c**) SART. (**d–j**) Self-report surveys. AC, attentional control; MW, mind wandering.

## Discussion

Multiple neurocognitive processes influence whether and how humans remember or forget, including preparatory processes that render certain moments more conducive to successful retrieval than others [5]. The present study provides further evidence that pupil-indexed suboptimal A-A states at retrieval are associated with impaired item recognition and source recollection [1, 6], and provides novel evidence that a real-time A-A intervention just prior to an attempt to remember can rescue item recognition memory. Because the trigger manipulation occurred exclusively at retrieval — following each encoding period — these results provide causal evidence that phasic reorienting during A-A-defined suboptimal states affects item recognition at the retrieval stage, independent of encoding quality. These findings illustrate how a closed-loop pupillometry paradigm with phasic reorienting during A-A suboptimal states can causally improve memory access, extending the readiness-to-remember framework from correlational observation to real-time intervention. We discuss the implications of the present findings for theory and application.

### Hippocampal and cortical retrieval mechanisms

The observed rescue of item memory but not source memory, combined with a credibly larger trigger effect on item than on source *d*^*′*^ (ER = 249.00), bears on theories of episodic retrieval. Dual-process models of recognition memory distinguish between familiarity — a rapid signal indexing item memory strength — and recollection — a slower, controlled process that recovers specific episodic details such as the encoding context. Extant data suggest that, relative to familiarity, recollection is more disrupted by division of attention at the time of retrieval [10, 84] and is differentially dependent on hippocampal pattern completion [85–87], with the latter being influenced by LC-NE/acetylcholine arousal and attentional modulatory input [17, 88–90]. Familiarity-based item recognition decisions can be affected by division of attention, though to a lesser extent than recollection [91–93], and differentially depend on medial temporal cortical computations [94–96]. While goal-directed attentional gain signals can increase familiarity [97–100], the influence of LC-NE arousal and attentional modulatory input on familiarity remains unclear.

The item recognition decisions in the present study may reflect familiarity-based memory in the absence of recollection or a mix of familiarity-based memory with and without non-criterial recollection (i.e., recollection of other encoding features beyond source details). The selective rescue of item memory, in the absence of a corresponding benefit for source memory, aligns with the dual-process view that familiarity and recollection rely on partially distinct neural substrates. We speculate that, in the current study, the reorienting trigger promoted an increase in arousal and a phasic reallocation of visuospatial attention to the location where the retrieval probe would appear, increasing neural gain and thus boosting the strength of medial temporal cortex familiarity elicited by the retrieval probe. Future studies are required to determine whether these effects are due to gain amplification of cortical signals that project to perirhinal and/or parahippocampal cortical structures from which familiarity signals emerge as well as whether such amplification is due to LC-NE modulatory effects or to effects of visuospatial attention.

By contrast, the phasic arousal and attentional reorienting elicited by the brief trigger was insufficient to increase the sustained, goal-directed attentional state needed to boost recollection-based memory decisions. It may be the case that a more sustained or semantically targeted reorienting cue is needed to facilitate recollection during suboptimal A-A states. Future work could test this prediction by comparing reorienting cues of varying duration, complexity, and/or semantic content — for example, reinstating the encoding goal cue itself — to determine whether richer interventions can additionally rescue source recollection.

### Intervening across the A-A state continuum

The present drift-corrected reclassification analysis offers a window into the heterogeneous nature of A-A suboptimal states and potential boundary conditions of real-time intervention. A key insight from the reclassification analysis is that the trials originally classified as SOS by the real-time algorithm fall along a continuum of A-A state deviation severity, ranging from threshold-proximal SOS — in which pupil-indexed arousal/attention deviates only slightly beyond the detection threshold — to deep SOS — in which states deviate substantially from the participant’s current physiological baseline, reflecting a more profound deviation from LC-NE tone optimality in the moment immediately preceding a retrieval trial.

This distinction provides an account of the pattern of results observed under reclassification. The conservative spline-correction procedure, by removing slow drift and recomputing thresholds against each block’s detrended distribution, effectively partitions the original SOS trials into these two categories. Trials that remain classified as SOS under both the real-time and drift-corrected methods represent deep SOS: A-A state deviations that are robust to the choice of baseline and detrending method.

Among these deep SOS, the trigger conferred a weaker and less certain memory benefit than among drift-driven SOS trials (ER = 7.03 vs. ER = 29.85), suggesting that when arousal/attention has deviated further from the optimal zone, a brief visual flickering cue may be insufficient to restore the preparatory state needed for successful memory retrieval.

By contrast, the approximately 55% of original SOS trials that were reclassified as OS by the drift-correction method represent threshold-proximal SOS — trials on which the real-time algorithm detected a threshold crossing, but the deviation was mild enough that, once slow drift is accounted for, these trials fall within the normal range of A-A state fluctuation. It is among these threshold-proximal SOS trials that the clearest trigger benefit emerged. Trials that received the reorienting cue showed credibly higher item memory than those that did not, with strong evidence (ER = 29.85). This pattern is consistent with the view that threshold-proximal A-A state deviations occupy a zone of recoverability; the arousal/attention system has drifted from optimal but has not disengaged or overengaged so deeply that a phasic reorienting signal cannot pull it back. Here, the brief visual trigger, by evoking a transient LC-NE-mediated phasic response and/or increase in visuospatial attention, appears sufficient to nudge the system from a mildly suboptimal state back toward the A-A threshold necessary for successful memory expression.

Taken together, this framing reinterprets the disappearance of the trigger benefit under drift-corrected reclassification not as evidence against the efficacy of real-time intervention, but as suggestive of a dose-response relationship between SOS severity and intervention efficacy. Indeed, the trigger produced directionally consistent benefits at both levels of reclassified suboptimality, and the pattern of simple effects — a stronger trigger benefit for drift-driven than drift-resistant SOS trials — is consistent with a fixed-perturbation account in which a brief A-A-boosting stimulus yields greater functional recovery when the distance to the inverted-U peak is smaller. However, the formal interaction contrasting these two trigger effects was inconclusive (ER = 2.27), and the present data lack the precision to definitively establish that the trigger operates differentially at different depths of suboptimality. The depth-of-suboptimality interpretation therefore rests on the converging pattern of simple effects, rather than on a credible interaction. Future studies with larger samples or adaptive designs that oversample deep SOS trials could provide the statistical precision needed to formally adjudicate this question.

Notably, although posterior mean item *d*^*′*^ was lower for drift-resistant than for drift-driven SOS trials in both trigger conditions, the comparison between drift-resistant and drift-driven untriggered trials did not yield a credible difference in *d*^*′*^ (ER = 2.32).

If drift correction captures a meaningful severity gradient, one might expect drift-resistant trials to show detectably lower *d*^*′*^ than drift-driven trials. Two considerations temper this concern. First, per-participant cell sizes for these conditions are modest, limiting sensitivity to detect what may be a small difference between neighboring points on the continuum. Second, the performance function may be relatively flat across the mildly-to-moderately suboptimal range and steepen only at the extremes — a pattern that would produce the observed result in which the broad reclassified-SOS vs. reclassified-OS contrast is large but the within-SOS gradient is undetectable at our sample size. Importantly, the depth interpretation does not rest on this comparison alone. It is also consistent with the lower *d*^*′*^ for drift-resistant than for drift-driven triggered trials (ER = 8.27), the differential trigger effects for drift-resistant and drift-driven SOS trials, and the constriction/dilation asymmetry discussed below.

### Intervening across low- and high-A-A states

In the planned test across all real-time SOS trials, the trigger credibly improved *d*^*′*^ for constriction trials (ER = 19.51) but not for dilation trials (ER = 2.20), with the interaction directionally favoring constrictions (ER = 3.92) though not reaching our criterion for credibility. The post hoc split showed the same pattern among drift-resistant SOS trials (i.e., the most deeply suboptimal states; ER = 24.53, 2.24, and 4.27). This pattern is consistent with the inverted-U framework’s prediction that hypoarousal states, situated on the ascending limb, are amenable to an A-A-boosting intervention that shifts the system toward optimality, whereas hyperarousal states, situated on the descending limb, should not benefit from, and may even be harmed by, additional arousal/attention. Among drift-driven SOS trials, by contrast, the evidence was anecdotal for both directions and for their difference (all ER *<* 3), which suggests that the differential effects of intervention may depend on whether suboptimality is far from versus closer to the peak of the inverted-U curve. While only suggestive, this pattern may inform future closed-loop interventions: Adaptive algorithms that detect the direction of the A-A departure could, in principle, deploy different intervention strategies for hypoarousal A-A versus hyperarousal A-A states depending on the degree of suboptimality — for instance, an alerting cue for extreme pupillary constrictions and a calming or refocusing cue for extreme pupillary dilations, whereas a common approach may suffice for state deviations closer to optimality.

### Pre-probe pupil, trait attention, and memory

Whereas the closed-loop design asks whether perturbing A-A state changes memory, the complementary trial-level analyses describe how memory varies with the A-A state assayed just before the retrieval probe. On triggered trials, larger pre-probe pupil predicted poorer item memory. If the reorienting cue raises arousal, the triggered trials that ended with the largest pupils may be those pushed past the inverted-U peak, including the dilation-defined SOS trials for which the trigger benefit was not credible.

Across participants, trait attention predicted item recognition, further supporting the readiness-to-remember framework [5, 6, 14]. Specifically, a latent factor combining task-based and self-reported attention predicted item memory, and participants in the top third of this factor recognized items with 0.44 *d*^*′*^ greater sensitivity than those in the bottom third (2.86 vs. 2.41). Because neither task-based nor self-reported indicators alone yielded a credible latent association, trait attention relevant to memory may be best captured by aggregating across multiple measurement methods, which are themselves only weakly intercorrelated. The relationship was weaker for source recollection and not credible. At the trial level, by contrast, SOS impaired source memory as well as item memory, with extreme evidence for both after reclassification. One possibility is that trait attention primarily affects the strength of item representations available at retrieval, whereas momentary A-A state additionally affects the controlled processes needed to recover encoding context.

### Limitations and future directions

Several limitations of the present study warrant consideration.

First, our real-time SOS detection relied on a fixed early-block baseline, which, as the spline-detrending analysis revealed, is susceptible to slow physiological drift. The reclassification analysis showed that the SOS impairment of item and source memory was robust to drift correction, but the overall trigger effect was not, and its clearest benefit appeared in drift-driven SOS trials, which the fixed baseline had flagged mainly because of drift. Future implementations of PRISM could incorporate adaptive, rolling baseline algorithms that are less sensitive to low-frequency drift while remaining causal.

Second, the reorienting trigger was a simple visual flicker — a low-level, non-semantic cue that may have boosted arousal, visuospatial attention, or both. While this design choice enabled experimental control by holding the cue constant across all triggered trials, it also limits mechanistic understanding of the trigger effect, as well as the ecological validity and potential potency of the intervention. Real-world applications might benefit from more targeted cues, such as brief reminders of the task goal or multi-sensory stimulation, which could potentially rescue both item and source memory.

Third, trigger efficacy varied widely across participants. Each participant’s trigger effect on item *d*^*′*^ (SOS trials with minus without trigger) ranged from *−* 1.37 to 1.43 (mean = 0.19, s.d. = 0.61), and 35 of 67 participants showed a positive effect. It is possible that the reorienting cue is most effective in individuals whose A-A fluctuations are typically small, such that their threshold crossings reflect recoverable departures. Testing this account will require designs that estimate each participant’s trigger benefit more precisely. To illustrate, by delivering more trigger and no-trigger trials per participant and/or by modeling the trigger effect hierarchically at the trial level.

Moreover, although the within-subject design controls for many individual differences, participants varied considerably in their SOS rates (52% to 90% of post-burn-in trials; mean = 68%) and trigger responsiveness. Further characterizing the individual-difference factors that moderate closed-loop efficacy will require larger samples and designs that estimate each participant’s trigger benefit with greater precision.

Fourth, while pupil size provides a continuous, non-invasive index of LC-NE-mediated arousal/attention, it is an indirect measure that reflects contributions from multiple autonomic and central nervous system pathways. The extent to which our findings generalize to other assays of A-A states, such as concurrent pupillometry-fMRI [e.g., using fMRI to index attentional states; 101] and/or pupillometry-EEG [e.g., using EEG posterior alpha to index attentional states; 6] paradigms, warrants further investigation.

Finally, the present study focuses on young adult participants with normal vision and no neurological disorders; the translational potential of PRISM for clinical populations with attentional dysregulation — such as individuals with ADHD, traumatic brain injury, and/or age-related cognitive decline — represents a promising but untested avenue.

### Conclusion

In summary, the present study demonstrates that real-time detection of and intervention on arousal-attentional suboptimal states immediately preceding attempts to remember can causally rescue item memory performance, providing closed-loop pupillometry evidence for the readiness-to-remember framework. The distinction between item and source memory rescue suggests that a brief reorienting cue is sufficient to boost familiarity-based recognition but not recollection of the encoding context. The spline-detrending reclassification suggests that the trigger’s efficacy depends on the depth and direction of suboptimality. The benefit was clearest for marginally suboptimal states within a zone of recoverability and for low-A-A states on the ascending limb of the inverted-U, although it was not credibly larger than for deeply suboptimal or high-A-A states. Together, these findings advance understanding of how moment-to-moment fluctuations in arousal and attention influence the expression of episodic memories, and they motivate real-time interventions tailored to both the severity and direction of arousal-attentional departure from optimality, with implications for educational practice, clinical intervention, and the design of adaptive cognitive technologies that seek to optimize human cognitive performance in real time.

## Materials and methods

### Participants

Seventy-nine participants were enrolled in the study (54 female participants; *mean age*=21.73 years, *s*.*d*.=1.92, range=18–25 years), based on sample sizes used in similar studies [6]. Participants were recruited from print and online advertisements at Stanford University and the surrounding community, were all native English speakers, reported not concurrently participating in any other study or in any study with contraindications to the primary hypotheses of the current study, reported no known neurological impairment, history of seizures and/or photosensitive epilepsy, nor severe dizziness, vertigo, and/or motion sickness, and did not possess an implanted pacemaker or defibrillator. All participants self-reported normal or corrected-to-normal vision, no known colorblindness, and normal or corrected-to-normal hearing in addition to passing a standardized vision test (ZEISS Online Vision Screening^2^), which included tests of visual acuity, contrast vision, color vision, and signs of astigmatism. All participants provided written informed consent and were compensated US$20 per hour, in accordance with procedures approved by Stanford’s Institutional Review Board. Data from 10 participants were excluded for the following reasons: (a) five participants were excluded from analyses owing to too few SOS trials; (b) another two participants for too many SOS trials (i.e., exceeding 2 s.d. of the mean proportion of SOS trials relative to all participants); (c) one participant was excluded for having source *d*^*′*^ less than or equal to zero; and (d) two participants were excluded from the study owing to failure to comply with task instructions. Two participants withdrew from the study altogether. Thus, data from 67 participants were retained for all analytics.

### Experimental design

Participants completed a 2.5-h session: set-up, vision test, and task instructions (30 min), goal-directed encoding and retrieval phases (1.5 h) and a separate individual differences assessment (30 min). Only the memory task included pupillometry measurements. Key components of the memory task (design, goal states, and common object pictures) were adapted from prior research that demonstrated the behavioral and functional neural correlates of item and source memory retrieval goals [6, 98]. The individual differences assessment was adapted from a prior study linking preparatory attention to item and source memory performance [6], and comprised two task-based assays of sustained attention — the gradual-onset continuous performance task (gradCPT) [39] and the sustained attention to response task (SART) [78] — as well as self-report questionnaires probing measures of media multitasking [79], attention-deficit/hyperactivity disorder (ADHD) [80], attentional impulsivity [81], attentional control over distraction and shifting [82], and spontaneous and deliberate mind wandering [82]. The individual-differences assessment was administered immediately after the real-time memory task and is reported in the Results herein.

### Goal-directed encoding and source memory retrieval task

All tasks were run using custom PsychoPy scripts in Python [102]. The fixation cross, goal cue, and object on each trial were presented centrally on a white background screen and were luminance- and chrominance-controlled and matched using the SHINE toolbox in MATLAB [103, 104]. Stimuli were presented on a Mitsubishi Diamond Pro 2070SB SuperBright Diamondtron cathode-ray tube display (June 2004 model) to ensure that low-level visual properties did not affect pupillary assays. The display was 40 cm wide × 30 cm tall at a resolution of 1024×768 px, where 400 px is approximately 15.6 cm. Participants’ eyes were situated approximately 50 cm from the monitor and performed the task in a stabilizing head- and chin-rest. A 5-point eye-tracker position calibration and validation procedure was performed prior to each block. All participants were run in the same testing environment — a dimly lit room in which the only light sources were the stimulus display and a dim floor lamp situated approximately 3 ft behind the participant; no fluorescent overhead lights were used during the session.

#### Encoding

Participants viewed 168 objects once across three intentional encoding blocks (7 min 56 s per block) of 56 trials each. Each encoding phase block preceded its corresponding retrieval phase block. On each trial (Fig 1a), a fixation cross appeared for 4.0 s, followed by one of two possible task orienting goal cues (‘pleasant or unpleasant’; ‘bigger or smaller’) for 1.6 s; a 0.1-s goal cue to stimulus interval was then followed by an object for 2.8 s. To minimize eye-gaze related pupillary artifacts, participants were instructed to keep their eyes fixated on the fixation cross until the goal cue or object appeared. Participants performed object classification based on a conceptual goal (does the object evoke feelings of pleasantness or unpleasantness) or a perceptual goal (is the object physically bigger or smaller in size on the screen, irrespective of real-world size), responding as quickly and as accurately as possible. Participants made each judgment with one of two button presses (pleasant or unpleasant; bigger or smaller) on a Sony PlayStation 5 DualSense^®^ Wireless Controller using their right thumb (square button to respond ‘pleasant’ or ‘bigger’, circle button to respond ‘unpleasant’ or ‘smaller’).

Participants received detailed verbal and visual instructions from the experimenter and were frequently given the opportunity to ask clarifying questions before moving on to the next instruction. There were eight interactive practice trials to ensure task comprehension before moving on to the real task blocks. Interactive practice was monitored closely by the experimenter and repeated until the participant demonstrated clear understanding of the task instructions and controller button-to-response mapping, typically occurring following the first or second round of practice trials.

In each encoding block, 28 objects were classified on the conceptual goal dimension (pleasant or unpleasant) and 28 on the perceptual dimension (bigger or smaller), yielding 84 conceptual object trials and 84 perceptual trials across blocks. Each object appeared either bigger (450×450 px) or smaller (150×150 px) in size on the screen, independent of goal cue, counterbalanced across participants through random assignment. Thus, objects were crossed in a 2×2 design, with 14 objects from each size crossing (‘pleasant or unpleasant?’ and bigger, ‘pleasant or unpleasant?’ and smaller, ‘bigger or smaller?’ and bigger, and ‘bigger or smaller?’ and smaller) appearing in a random order per block (with the constraint that a particular goal — for example, ‘pleasant or unpleasant’ — could not appear more than four times consecutively). Thus, each participant received a random assignment of 168 goal-object pairings at encoding.

#### Retrieval

Following each encoding block, all participants were given a 5-min interphase rest period and were permitted to stand up, stretch, walk around the dimly lit room, and/or sit back and close their eyes. During this interphase retention interval, participants were also played a medley of relaxing music. Following the interphase period, participants performed a retrieval phase for objects from that immediately preceding encoding block. Specifically, at retrieval, they viewed 84 objects — 56 studied and 28 novel foils — across each of three goal-directed retrieval blocks of 84 trials each (252 total objects). Similar to the encoding phase, objects appeared in a random order per block (with the constraint that an object’s associated encoding phase goal — for example, ‘pleasant or unpleasant’ — could not appear more than three times consecutively).

Participants were instructed that in each retrieval block, medium-sized versions of objects from the immediately preceding encoding phase block would be intermixed with new objects that they have never seen before in the experiment. On each retrieval trial, participants made a four-alternative forced-choice (4-AFC) memory judgment on each medium-sized test object (300×300 px), wherein they decided whether they: (1) remember having performed the size decision task (i.e., bigger or smaller before) with the object at encoding, (2) remember having performed the pleasantness decision task (i.e., pleasant or unpleasant before) with the object at encoding, (3) remember having encountered the object at encoding, but cannot remember which decision task was performed, or (4) do not remember having encountered the object and thus perceive it as new.

Each retrieval trial (Fig 1c) began with a 4-s intertrial interval (ITI) consisting of a fixation cross. The final 2 s of this interval — for the first 20 trials of each retrieval block — constituted the relevant analytic window for “burn-in” trials to establish robust baseline pupillometry metrics before threshold-based assessment of A-A states and, on a subset of trials, triggering commenced. Beginning on trial 21, following the 4-s ITI, real-time pupillometry was conducted across either one or two 1-s real-time sampling windows to assess momentary pupil dilation (note that if a SOS was not detected in the first 1-s window the second 1-s window was assessed, which provided more opportunities for detection of a SOS and then, on a subset of trials, real-time triggering). A second 1-s sampling window also appeared when the first 1-s window contained excessive missing pupil data (pre-determined to be *≥* 20% of pupil samples). Data from each real-time window then underwent real-time blink padding (100 ms) and linear interpolation; if more than 50% of the real-time time series following blink padding were missing values, the real-time window sample was rejected. Moreover, we implemented a real-time trial recovery approach to increase power by reducing the number of trials with no reliable real-time triggering data; here, we “recycled” trials if both 1-s real-time sampling windows were impacted by excessive missing pupil data. Recycling consisted of displaying a control circle (luminance-matched and gray-tan, with a radius of 150 px, the color of which was determined by averaging the RGB values for all possible object probe stimuli) in lieu of an object probe, and the trial was restarted for up to three additional repetitions (totaling up to four possible trial initiations per object) before marking real-time data for that object unusable. Participants were instructed to not respond in the event they received a circle trial, but were not told the purpose of these circles until session debriefing.

If the mean pupil assay during the 1-s real-time sampling window exceeded a threshold of *±* 1 s.d. from the fixed baseline mean (computed once across the 20 “burn-in” baseline trials), a reorienting cue could be triggered (Fig 1c). The reorienting cue consisted of a centrally presented, flickering circle (200-ms flicker period; 75 px radius) presented for 416 ms. To ensure balanced delivery, reorienting cues were delivered on 50% of threshold-crossing trials, with the remaining 50% receiving an equivalent-duration fixation cross. Assignment was balanced within each successive pair of SOS trials in a block: when the block’s trigger and no-trigger counts were equal, a coin flip chose the condition, and the next SOS trial received the other one.

Following the trigger decision period, a 1-s post-trigger fixation appeared immediately before the object probe was presented. Similar to the encoding phase blocks, participants were instructed to keep their eyes fixated on the fixation cross until the object probe appeared to minimize eye-gaze related pupillary artifacts. Object probes remained on screen for up to 5 s or until response, with participants instructed to respond as quickly and as accurately as possible.

Participants made each 4-AFC memory judgment (pleasant or unpleasant before; bigger or smaller before; old; or new) with one of four button presses on a Sony PlayStation 5 DualSense^®^ Wireless Controller using their right thumb (square button to respond ‘bigger/smaller before’, circle button to respond ‘pleasant/unpleasant before’, triangle to indicate remembering having encountered the object but not remembering which decision was made, and the × button to indicate that the object is new). Prior to the retrieval task, participants received detailed verbal and visual instructions from the experimenter and were given the opportunity to ask clarifying questions. There were eight interactive practice trials designed to ensure task comprehension; interactive practice was monitored closely by the experimenter and repeated until the participant demonstrated clear understanding of the task instructions and controller button-to-response mapping, which typically occurred following the first or second round of practice trials. To accurately indicate whether an object had been perceptually or conceptually encoded requires object/item recognition and recollection of how the object was processed at encoding [6]. To accurately classify an object as “old” but without source recollection requires recognition based on item familiarity with or without non-criterial recollection. Accurate novelty detection was likely based on weak item familiarity.

### Individual differences assay

#### Gradual-onset continuous performance task (gradCPT)

In this task-based assay of sustained attention [39], participants viewed a continuous stream of round, grayscale city and mountain scenes (300×300 px, equated for mean luminance) that gradually transitioned from one image to the next. They were instructed to press the space bar for city scenes (frequent; 90% of trials) and to withhold responding for mountain scenes (infrequent; 10% of trials). The task comprised 500 trials over 10 min. Each image transition spanned 24 frames at 50 ms per frame (1.2 s per trial), and the same scene could not appear on consecutive trials. The stream began with a transition from scrambled noise into the first scene and ended with a transition back to noise; this final transition was not scored, leaving 499 trials per participant, of which 34–67 were mountain scenes (*M* = 50.7; 10.2%).

#### sustained attention to response task (SART)

In this task-based assay of sustained attention [78], participants viewed a continuous stream of displays, each showing six colored shapes in a ring around a central fixation dot. On each display, the six shapes were either all circles or all squares. One shape type was frequent (540 of 600 trials, 90%) and the other infrequent (60 trials, 10%). Participants pressed one key (‘1’ or ‘2’) for each shape type on every display, as quickly and as accurately as possible. Which shape was infrequent and which key went with each shape were counterbalanced across participants. Each display stayed on screen for 800 ms, and responses were recorded only during that time. The next display followed immediately, with no blank interval between displays, so the 600 trials, presented in random order, took 8 min. Shape colors changed from display to display. The fixation dot turned from black to white when a key press registered, and a reminder of the key mapping stayed at the bottom of the screen throughout. Only the first key press on each display counted, and displays without a response were scored as errors. A 50-trial practice block, with the same 10% rate of infrequent shapes, came before the main task.

#### Questionnaires

A modified Media Multitasking Inventory (MMI) [79] assessed the hours per week spent on each of eight media activities: reading; homework other than reading; listening to music or other audio; playing video games; browsing the internet; texting, social media, or instant messaging; talking on the phone or video chatting; and other computer activities. Participants rated, once per pair, how often they engaged in each pair of activities simultaneously on a four-point scale.

ADHD symptoms were assessed with Part A of the Adult ADHD Self-Report Scale (ASRS) [80], six items rated on a five-point frequency scale. Impulsivity was assessed with the Barratt Impulsiveness Scale-11 (BIS-11) [81], 30 items rated on a four-point scale and organized into attentional, motor, and non-planning second-order subscales. Everyday attentional control was assessed with two four-item subscales measuring attentional distractibility (AC-D) and difficulty shifting attention (AC-S) [82]. Deliberate (MW-D) and spontaneous (MW-S) mind wandering were each assessed with four items rated on seven-point scales [82]. The MMI and BIS-11 each contained an embedded attention-check item.

#### Latent attention factor

Trait attention was modeled with a Bayesian one-factor confirmatory factor analysis of ten indicators, one per construct: MMI, gradCPT *d*^*′*^, gradCPT RT CoV, SART rare-shape accuracy, and the ASRS, BIS-11 attentional, AC-D, AC-S, MW-S, and MW-D scores. To avoid counting a construct twice, we omitted scores that overlapped with a retained indicator: the gradCPT commission and omission error rates, from which *d*^*′*^ is computed, and the BIS-11 total, which includes the attentional subscale. We also omitted scores that mainly index something other than attention: the BIS-11 motor and non-planning subscales, which measure other forms of impulsivity, and the SART RTs and frequent-shape accuracy, which mainly index response speed. Each indicator was *z*-scored and signed so that higher values indicate better attention (the MMI, ASRS, BIS-11, AC-D, AC-S, MW-D, MW-S, and RT CoV scores were reversed).

Let *z*_*ij*_ denote the signed *z*-score of participant *i* (*i* = 1, … , 67) on indicator *j* (*j* = 1, … , 10). Indicator scores were modeled as *z*_*ij*_ *∼* Normal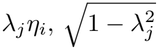, where *λ*_*j*_ is the loading of indicator *j* and *η*_*i*_ *∼* Normal(0, 1) is the factor score of participant *i*, so that each loading *λ*_*j*_ is the standardized loading (i.e., the indicator’s correlation with the factor). Loadings were given the prior *λ*_*j*_ = 2*·*Beta(2, 2) *−* 1, which is weakly informative on (1, 1). Only observed indicator values entered the likelihood, so participants missing one assessment contributed through their remaining indicators. Posterior draws were sign-aligned after sampling by flipping draws whose loadings summed to less than zero; convergence diagnostics were computed on the aligned draws. Composite reliability was summarized with McDonald’s *ω* [105], computed on every posterior draw as 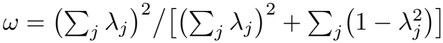 , and factor scores were each participant’s posterior mean of *η*_*i*_.

We related the factor to memory in three complementary ways. First, we related factor scores to item and source memory *d*^*′*^ with the Bayesian regression described below. Second, in a structural extension of the model, we added standardized *d*^*′*^ as Normal 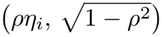 with *ρ∼* Uniform(*−* 1, 1), fit separately for each outcome, to estimate the latent correlation *ρ* while accounting for uncertainty in the factor scores. Third, as a model-free check, we averaged each participant’s available signed *z*-scores into a unit-weighted composite and related it to item and source memory *d*^*′*^ in the same way. Because the composite weights all indicators equally, it tests whether the factor’s association with memory depends on the estimated loadings. To describe the size of the association, we also split participants into thirds of the factor-score distribution (equal-count tertiles) and compared mean item *d*^*′*^ between the top and bottom thirds. For the factor scores, the composite, and each indicator separately (Fig 7), we regressed *z*-scored *d*^*′*^ on the *z*-scored predictor with the default priors of brms [83], including a flat prior on the slope, so the standardized slope *β* is on the correlation scale; we report its posterior mean and 95% HDI.^3^

### Pupillometry data acquisition

During encoding and retrieval, pupillary data were recorded in a dark room where the only source of light was the stimulus display monitor and a dimly lit floor lamp. Real-time eye-tracking was monitored on the EyeLink Host PC by a researcher positioned behind a folding room divider. Pupillary data were recorded from the right eye using an EyeLink 1000 Eye Tracker system (SR Research^4^) at a sampling rate of 1000 Hz. Participants were seated approximately 50 cm from the eye-tracker and stimulus monitor with a stabilizing head- and chin-rest. Five-point eye-tracking calibration and validation steps were completed before each encoding and retrieval block. Trial-level pupillary and behavioral data (response and response time) were synced using a custom Python script with event message tags.

### Data analyses

Trial-level pupillometry and individual-differences analyses were conducted in Python (v3.14) [107] within a marimo (v0.25) reactive notebook [108], using pandas (v3.0) [109], NumPy (v2.4) [110], SciPy (v1.18) [111], PyMC (v6.3) [112] with the nutpie (v0.16) No-U-Turn sampler [113–115], ArviZ (v1.3) [116], matplotlib (v3.11) [117], and seaborn (v0.13) [118]. All Bayesian models used the same sampler settings (four chains of 4,000 iterations with 1,000 warmup, yielding 12,000 posterior draws; random seed 512). Convergence was assessed with rank-normalized 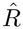 and bulk and tail effective sample sizes [119]. Highest-density intervals (HDIs) were computed as the narrowest interval containing the stated posterior mass.

#### Memory behavioral data analyses

Behavioral analyses focused on metrics of memory retrieval accuracy. For studied objects, each 4-AFC response was classified at two levels. At the item level, a response was classified as an item hit if the participant endorsed the object as previously encountered — i.e., responded “size before”, “pleasantness before”, or “old” — and as an item miss if the participant responded “new”. At the source level, a response was classified as a source hit if the participant correctly identified the encoding task (e.g., responded “size before” to an object encoded under the size goal) and as a source miss otherwise (i.e., attributed the wrong encoding task, responded “old” without source specification, or responded “new”). For novel objects, a response was classified as an item false alarm if the participant endorsed the object as old (responded “size before”, “pleasantness before”, or “old”), and as a source false alarm if the participant specifically attributed an encoding source (responded “size before” or “pleasantness before”).

We adopted signal-detection theory [120] to compute log-linear corrected *d*^*′*^ statistics (*Z*_hit_ *− Z*_false alarm_) for each A-A state crossed with trigger condition (Fig 1b) — i.e., *d*^*′*^ statistics were computed separately for the OS, SOS without trigger, SOS with trigger conditions. When computing item memory *d*^*′*^, hit rate was calculated as the proportion of old items (i.e., encountered during the encoding phase) that elicited a “size before”, “pleasantness before”, or “old” response by the participant; false-alarm rate was calculated as the proportion of novel foils that were incorrectly responded to with a “size before”, “pleasantness before”, or “old” response by the participant. For source memory *d*^*′*^, hit rate was calculated as the proportion of old items on which a participant correctly retrieved the source (i.e., correctly indicated the classification judgment performed on the object during the encoding phase); false-alarm rate was calculated as the proportion of novel foils that were incorrectly given a “size before” or “pleasantness before” response by the participant.

Performance outliers were identified as any participant with item memory *d*^*′*^ or source memory *d*^*′*^ less than or equal to 0 and/or any participant who had normatively few or many SOS trials (i.e., more than 2 s.d. from the mean proportion of SOS trials relative to all participants). Data from an outlier were excluded from all analyses.

We estimated condition means for item and source memory *d*^*′*^ with Bayesian linear mixed models in a cell-means parameterization, *d*^*′*^ *∼*0 + condition + (1 | participant), so each population-level coefficient is one condition’s mean. Each participant contributed one *d*^*′*^ per condition, and these values were correlated across conditions (*r* = 0.61–0.69 for the real-time conditions).^5^ Each model had a Gaussian likelihood. We wrote the models in PyMC [112] using the default priors of brms [83]: each condition mean had an improper flat prior, and the by-participant and residual standard deviations had half-Student-*t*(3, 0, 2.5) priors, with the participant effects non-centered. We sampled each posterior with the No-U-Turn Sampler in nutpie [113–115]. Each model ran four chains of 4,000 iterations, and we discarded the first 1,000 of each chain as warmup, leaving 12,000 posterior draws. Every parameter had 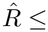 1.01 and a bulk effective sample size above 1,500, and no model had a divergent transition. We tested three directional hypotheses: SOS *<* OS, SOS with trigger *>* SOS, and SOS with trigger *<* OS. For each condition we report the posterior median (*m*) and its 95% highest-density interval (HDI). For each hypothesis we computed the contrast *b*, the difference between the two condition means, on every posterior draw. We report its posterior mean and its 90% equal-tailed credible interval (CrI). Following `hypothesis()` in brms, we measured the evidence for each hypothesis with its evidence ratio (ER), the posterior probability of the predicted direction divided by the posterior probability of the opposite direction. For SOS *<* OS, for example, ER = *P* (*b <* 0 |data)*/P* (*b >* 0 |data).^6^ The ER is therefore the Bayes factor for the hypothesis against its opposite direction.^7^ We considered a hypothesis supported when its 90% CrI lay entirely on the predicted side of zero.^8^ Following Lee and Wagenmakers [121], we describe ERs of 1–3, 3–10, 10–30, 30–100, and above 100 as *anecdotal, moderate, strong, very strong*, and *extreme* evidence.

To test whether the trigger effect differed between item and source memory, we computed for each participant *i* the difference between their item and source trigger effects, 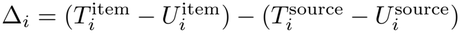, where *T* and *U* are *d*^*′*^ on SOS trials with and without trigger. Because both trigger effects come from the same trials, this difference is taken within participant. We modeled Δ_*i*_ *∼* Normal(*μ, σ*) with an improper flat prior on *μ* and a half-Student-t(3, 0, 2.5) prior on *σ*, sampled as above, and tested *μ >* 0 with its ER.

#### Individual differences scoring

For the gradCPT, we computed four performance indices. The commission error rate was the proportion of mountain trials with a response, and the omission error rate was the proportion of city trials without a response. Sensitivity (*d*^*′*^) was computed with the log-linear correction implemented in the R package psycho [122], treating correctly withheld mountain trials as hits and withheld city trials as false alarms, such that higher *d*^*′*^ indicates better discrimination of when to withhold a response. This coding gives the same *d*^*′*^ as the conventional coding (responses to city trials as hits, responses to mountain trials as false alarms), because each of its rates is the complement of a conventional rate and *z*(1 *−p*) = *−z*(*p*), with or without the log-linear correction.

Finally, we computed mean response time and its coefficient of variation (RT CoV; 100 × s.d./mean) across all trials with a valid response. The number of mountain trials, which ranged from 34 to 67 across participants, was not credibly related to any of these measures or to the latent attention factor scores described below. In Bayesian regressions of each *z*-scored measure on the *z*-scored number of mountain trials, every 95% HDI of the standardized slope included zero (*β* from *−* 0.22 to 0.16), and two-sided Bayes factors from the default Bayesian Pearson test favored no relation (BF_10_ = 0.18–0.69; *n* = 66).

For the SART, we computed accuracy, the proportion of displays with the correct key press, separately for frequent-shape displays and for infrequent-shape displays (rare-shape accuracy), as well as the median RT for correct frequent-shape trials. Displays without a response counted as errors. Participants failing to respond on *≥* 20% of trials were to be excluded; however, none met this criterion (missed responses: *mean*=3.2%, *maximum*=16.7%).

For the MMI, pairwise co-use ratings were coded 0, 0.33, 0.67, and 1, and summed for each medium across its pairings with every other medium (*m*_*i*_). Each participant’s MMI was the hours-weighted mean of these sums, ∑_*i*_(*h*_*i*_ *×m*_*i*_)*/* ∑_*i*_ *h*_*i*_, where *h*_*i*_ is the hours per week spent on medium *i*. Item scores were averaged for the ASRS (recoded 0-4), AC-D, AC-S, MW-D, and MW-S. For the BIS-11, 11 items were reverse-scored and items were summed into total, attentional (8 items), motor (11 items), and non-planning (11 items) scores. For all questionnaires, higher scores indicate more of the named construct (e.g., more inattention, impulsivity, or mind wandering).

Measures were treated as missing for the participant without gradCPT and SART data (*n* = 1), the participant without questionnaire data (*n* = 1), the three participants with invalid gradCPT response times (RT measures only), and the two participants who failed the MMI attention check (MMI only). No participant failed the BIS-11 attention check or met the SART exclusion criterion. Analyses used all available data for each pair of measures (*n* = 63–67).

#### Participant-level memory sensitivity for individual-differences analyses

For analyses relating memory to individual differences in trait-level attention, item and source memory *d*^*′*^ were computed for each participant by pooling all untriggered trials (OS and SOS without trigger), using the trial classifications, response coding, and log-linear correction described above. Untriggered trials were used so that trait-level memory was not influenced by the intervention (item *d*^*′*^: *mean* = 2.60, *s*.*d*. = 0.70; source *d*^*′*^: *mean* = 2.22, *s*.*d*. = 0.70; *n* = 67).

#### Trial-level pupillometry data analyses

Pupillometry data were preprocessed using the eyeris glassbox protocol with default parameters [77]. Specifically, the 1000-Hz data were first deblinked using a 50-ms symmetrical NaN-padding procedure around each missing sample. Then, physiologically unlikely samples were removed on the basis of speed (i.e., median absolute deviation (MAD) computed against a threshold from sample-to-sample) [123]. Missing pupil samples with a max gap *≤* 250 ms were then filled-in using linear interpolation, followed by lowpass filtering of the time series for smoothing (end of passband frequency: 4 Hz, start of stopband frequency: 8 Hz, required maximal ripple within passband: 1 dB, required minimal attenuation within stopband: 35 dB). Finally, all trial-level epochs were *z*-scored within a block, within each participant, rather than across blocks, to account for time-on-task effects. Burn-in trials, trials without a response, and control-circle (“recycled”) trials were excluded, as were epochs rejected during visual inspection of the eyeris quality-control reports.

Each trial’s pre-probe pupil was the mean of the block-wise *z*-scored pupil over its 1-s post-trigger (pre-stimulus) fixation epoch. For studied objects, item and source memory outcomes were coded as hits or misses following the definitions in **Memory behavioral data analyses**. We tested trial-level relationships with Bayesian logistic mixed-effects models written in PyMC [112], with a Bernoulli likelihood and logit link. Each studied-object trial contributed one observation for item memory and one for source memory. The fixed effects were hit *∼*pupil *×* memory, with memory sum-coded (item = *−* 0.5, source = +0.5), so the pupil term is the average slope across memory types and the interaction is the source minus item memory difference in slopes. The model included uncorrelated by-participant random intercepts and random slopes for pupil and memory, (1 + pupil + memory || participant). Because an item miss is necessarily also a source miss, a trial’s two observations are not independent; rather than model this dependence with a by-trial random intercept, we also fit hit *∼* pupil models separately for item and source memory, in which each trial contributes one observation. Each separate model had correlated by-participant random intercepts and pupil slopes, (1 + pupil | participant).

All models used the default priors of brms [83]. Each population-level slope had an improper flat prior. The intercept had a Student-t(3, 0, 2.5) prior, placed on the intercept with the predictors mean-centered. Each by-participant standard deviation had a half-Student-t(3, 0, 2.5) prior, and the intercept-slope correlation in the separate models had an LKJ(1) prior, which for two random terms is uniform on (-1, 1).

By-participant effects were non-centered. We sampled each posterior with the No-U-Turn Sampler in nutpie [113–115]. Each model ran 4 chains of 4,000 iterations, and we discarded the first 1,000 of each chain as warmup, leaving 12,000 posterior draws per model. Every pupil slope had 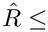 (maximum 1.001) and a bulk effective sample size of at least 3,249, and no model had a divergent transition.

On each posterior draw, we computed the item slope as the average slope - 0.5 *×* the interaction and the source slope as the average slope + 0.5 *×* the interaction. For each slope we report the posterior mean and its 90% HDI. As for the *d*^*′*^ models, we measured the evidence for a negative slope with the evidence ratio ER = *P* (*b <* 0 |data)*/P* (*b >* 0 |data), where values below 1 favor a positive slope.^9^ We fit the models to SOS trials with trigger.

### Offline spline-detrending analysis

To validate the robustness of PRISM’s real-time SOS classification and to operationalize a continuum of A-A state suboptimality, we implemented an offline spline-detrending analysis that reclassifies each trial’s A-A state independent of slow pupil drift. The procedure was applied separately to each participant and each of the three retrieval blocks.

#### Time series construction

For each block, we constructed a continuous time series by concatenating the preprocessed pupil data from the real-time monitoring windows across all trials in trial order. Inter-trial gaps — corresponding to the pre-stimulus ITI (4 s), trigger or equivalent fixation period (0.4 s), post-trigger fixation (1 s), and stimulus presentation period (5 s) — were padded with NaN values (10.4 s per trial at 1000 Hz) to preserve the temporal structure of the block while excluding non-real-time-monitoring-window data from the spline fit.

#### Spline fitting

A natural cubic spline with five degrees of freedom was fit to the padded time series using only the non-NaN samples from the real-time monitoring windows, capturing slow non-stationary trends in the pupil signal including linear drift and exponential decay components characteristic of time-on-task effects. The fitted spline was then evaluated at all timepoints — including the NaN-padded intervals — yielding a continuous drift estimate across the full block.

#### Trial-level drift correction

For each trial, we computed the mean of the spline-predicted values during that trial’s real-time monitoring window, representing the estimated drift level at the time of that trial. Each trial’s monitoring-window pupil size was then corrected by subtracting this trial-specific spline mean.

#### Reclassification

Unlike the real-time algorithm, which computed thresholds from a fixed baseline initialized over the first 20 “burn-in” trials, reclassification thresholds were derived from the global distribution of spline-detrended pupil values across all real-time monitoring samples within the block. Trials were reclassified as SOS if the drift-corrected monitoring-window mean exceeded ±1 s.d. of the global detrended mean, and as OS otherwise. This procedure leverages information from the entire block — including trials that were unavailable to the real-time classifier — providing a more stable reference distribution for threshold computation.

#### Six-cell condition matrix

Crossing the original real-time classifications (OS, SOS without trigger, SOS with trigger) with the drift-corrected reclassifications yielded a six-cell condition matrix (Table 1). Trials originally classified as SOS that retained their SOS classification after drift correction were termed “drift-resistant SOS” trials, representing A-A state departures severe enough that removing the drift component did not alter their classification. Trials originally classified as SOS that were reclassified as OS were termed “drift-driven SOS” trials, indicating that their original SOS classification was attributable primarily to drift rather than a substantial A-A state departure. This crossing enabled post hoc testing of whether trigger efficacy varies with the inferred depth and direction of suboptimality along the A-A state continuum.

### Asymmetry and post hoc models

The planned test of the asymmetry hypothesis compared the trigger effect between constriction- and dilation-defined SOS trials across all real-time SOS trials. Post hoc, we analyzed the reclassified conditions (i.e., the six cells) and split the constriction and dilation comparison by drift-resistant and drift-driven SOS trials. All of these analyses used the same models, including the by-participant random intercept, and the same directional tests as the real-time conditions.

## Data availability

Data are publicly available via the Open Science Framework with identifier 4epcn (https://osf.io/4epcn).

## Code availability

Analytic and custom PRISM task code for this study are publicly available via the Open Science Framework with identifier 4epcn (https://osf.io/4epcn) and on GitHub (https://github.com/shawntz/prism).

## Acknowledgments

We are grateful to T. Bonnen, T. Coleman, S. Delp, J. Fan, A. Franco, K. Grill-Spector, M. Harrison, B. Levy, D. Miller, R. Poldrack, Subbulakshmi S., E. Traister, A. Turnbull, and A. Xue for helpful comments and suggestions throughout the lifecycle of this project. This research was supported by the Wu Tsai Human Performance Alliance Agility Project Grant (to A.D.W., K.P.M., and S.T.S.), the Stanford Wu Tsai Neurosciences Institute’s Center for Mind, Brain, Computation and Technology (to S.T.S.), and a Stanford University Ric Weiland Graduate Fellowship in the Humanities & Sciences (to S.T.S.).

## Footnotes

1 This comparison and the drift-resistant comparison in the next section (ER = 7.03) use the same 2,037 triggered trials but different untriggered trials. Here, SOS without trigger includes every untriggered trial classified as SOS after drift correction: the 2,032 real-time SOS trials that remained SOS and 527 real-time OS trials reclassified as SOS (Table 1). Because the real-time algorithm never flagged those 527 trials, they could not have been triggered, and thus the drift-resistant comparison leaves them out.

2 https://visionscreening.zeiss.com/en-US

3 A flat prior leaves a Bayes factor against *β* = 0 undefined, so Bayes factors came from the default Bayesian Pearson correlation test [106]: one-sided BF_+0_ or BF_*−*0_ with *ρ* uniform over the predicted half, and two-sided BF_10_ with *ρ* uniform on (*−* 1, 1) where no direction was predicted.

4 https://www.sr-research.com

5 The by-participant random intercept accounts for this correlation, so contrasts between conditions are estimated within participants.

6 Posterior odds equal the Bayes factor times the prior odds. Because the condition means have flat priors, every contrast between them also has a flat prior, which gives the two directions equal prior probability.

7 It is not a Bayes factor against a point null (*b* = 0), which is undefined under improper priors, and it is on a different scale from the BF_+0_ values we report for the individual-differences correlations.

8 For a one-sided hypothesis this is the same as a posterior probability above 0.95, or an ER above 19.

9 The slopes have flat priors, so this ER is the Bayes factor for a negative slope against a positive one.

